# The PRR14 heterochromatin tether encodes modular domains that mediate and regulate nuclear lamina targeting

**DOI:** 10.1101/788356

**Authors:** Kelly L. Dunlevy, Valentina Medvedeva, Jade E. Wilson, Mohammed Hoque, Trinity Pellegrin, Adam Maynard, Madison M. Kremp, Jason S. Wasserman, Andrey Poleshko, Richard A. Katz

## Abstract

A large fraction of epigenetically silent heterochromatin is anchored to the nuclear periphery via “tethering proteins” that function to bridge heterochromatin and the nuclear membrane or nuclear lamina. We identified previously a human tethering protein, PRR14, that binds heterochromatin through an N-terminal domain, but the mechanism and regulation of nuclear lamina association remained to be investigated. Here we identify a centrally located, evolutionarily conserved PRR14 nuclear lamina binding domain (LBD) that is both necessary and sufficient for positioning of PRR14 at the nuclear lamina. We also show that PRR14 associates dynamically with the nuclear lamina, and provide evidence that such dynamics are regulated through phosphorylation of the LBD. We also show that the evolutionary conserved PRR14 C-terminal Tantalus domain encodes a PP2A phosphatase recognition site that regulates PRR14 nuclear lamina association. The overall findings demonstrate a heterochromatin anchoring mechanism whereby the PRR14 tether simultaneously binds heterochromatin and the nuclear lamina through two modular domains. Furthermore, the identification of a modular LBD may provide an engineering strategy for delivery of cargo to the nuclear lamina.

## Introduction

The eukaryotic DNA genome is organized into chromatin and is enclosed by the nuclear envelope that forms the border of the nuclear organelle. The nuclear envelope is a double membrane (Buchwalter et al., 2019; Hetzer, 2010; Ungricht and Kutay, 2015) that in multicellular organisms, includes the nuclear lamina protein framework that lies underneath the inner nuclear membrane (Dittmer and Misteli, 2011; Gruenbaum and Foisner, 2015). The nuclear lamina is composed of the intermediate filament proteins Lamin A/C, Lamin B1 and Lamin B2. In addition to acting as a nuclear framework, the nuclear lamina serves as a docking site for heterochromatin, which forms a characteristic silent “peripheral heterochromatin compartment” decorated with the repressive heterochromatic histone tail modifications, histone 3 lysine 9 di- and tri-methyl (H3K9me2 and H3K9me3) (Buchwalter et al., 2019; Gordon et al., 2015; Gruenbaum and Foisner, 2015; Lemaitre and Bickmore, 2015; Meister and Taddei, 2013; Padeken and Heun, 2014; Politz et al., 2013; Shevelyov and Nurminsky, 2012; Shevelyov and Ulianov, 2019; van Steensel and Belmont, 2017; Wong and Reddy, 2015). This heterochromatin compartment positioned at the nuclear lamina is the focus of much interest, as it provides a critical function in housing lineage-inappropriate repressed genes and gene-poor DNA (Becker et al., 2016; Peric-Hupkes et al., 2010; Reddy et al., 2008; Stancheva and Schirmer, 2014; Zhou et al., 2011). Furthermore, perturbations of heterochromatin, or gene positioning at the nuclear periphery, may contribute to disease and aging (Bank and Gruenbaum, 2011; Butin-Israeli et al., 2012; Davidson and Lammerding, 2014; Dittmer and Misteli, 2011). Although there has been significant progress in mapping local and global 3D chromatin organization (Dekker et al., 2017), an understanding of the mechanisms underlying such organization is limited. Regarding peripheral heterochromatin organization, historical and emerging findings, including our own, have shown that proteins function as “tethers” to attach heterochromatin to the nuclear periphery.

Early studies identified the Lamin B Receptor (LBR) (Olins et al., 2010; Ye and Worman, 1996) as a multi-pass inner nuclear membrane protein that binds to peripheral H3K9me3 heterochromatin through the prolific adapter protein, heterochromatin protein 1 (HP1) (Canzio et al., 2014; Lomberk et al., 2006; Zeng et al., 2010). The HP1 bivalent adapter function is mediated by an N-terminal chromodomain (CD) that is a ligand for H3K9me2/3 and a C-terminal chromoshadow domain (CSD) that recruits numerous <u>P</u>X<u>V</u>X<u>L</u> motif-containing partners to heterochromatin (Lechner et al., 2005; Machida et al., 2018; Nozawa et al., 2010). Using a cell-based epigenetic silencing factor screen, we identified previously an unstudied, widely expressed, 585 amino acid human protein, Proline Rich 14 (PRR14) (Poleshko and Katz, 2014; Poleshko et al., 2014; Poleshko et al., 2013). We showed that PRR14 localized strongly to the nuclear lamina, and data mining revealed that PRR14 had been detected as a binding partner of HP1 in two independent screens (Nozawa et al., 2010; Rual et al., 2005). We also determined previously that PRR14 encodes a modular N-terminal heterochromatin domain (1-135) containing a functional HP1-partner <u>P</u>X<u>V</u>X<u>L</u> motif (<u>L</u>A<u>V</u>V<u>L</u>) at positions 53-57. We hypothesized that PRR14 could function as a tether to position HP1-bound H3K9me3 heterochromatin at the nuclear lamina, and PRR14 knockdown experiments demonstrated such a role (Poleshko et al., 2013).

LBR and PRR14 tethering proteins are now both characterized as associating strongly with the nuclear periphery, and binding to and constraining adjacent heterochromatin (Shevelyov and Ulianov, 2019). The tethering mechanism implicates a simple bivalent attachment mechanism, through the HP1 adapter molecule. The identification of modular tether domains that can independently bind the nuclear periphery and heterochromatin can provide compelling confirmation of tether function.

We found that unlike LBR, PRR14 does not associate with nuclear membranes, but is instead dependent on the nuclear lamina component Lamin A/C for localization to the nuclear periphery (Poleshko et al., 2013). Also unlike LBR, which remains membrane-associated during mitosis, PRR14 behaves as a soluble heterochromatin binding protein. For example, PRR14 was found to disperse in metaphase, and colocalize with HP1 on chromosomes immediately at the onset of anaphase (Poleshko et al., 2013). We proposed that in addition to tethering, PRR14 may thereby have a mitotic “bookmarking” role for specification of heterochromatin for return to the nuclear lamina at the end of mitosis (Poleshko and Katz, 2014). Overall, our findings revealed that PRR14 binds HP1-heterochromatin similarly to LBR, but the nonmembrane-binding character may provide novel functions.

Recent studies identified the *C. elegans* CEC-4 protein as a membrane-associated heterochromatin tether. CEC-4 encodes an HP1-like CD that interacts directly with H3K9me1/2/3, therefore obviating the need for an HP1 adapter (Gonzalez-Sandoval et al., 2015). The identification of CEC-4 indicates that tethering through the H3K9me-modifications as anchoring points is conserved through evolution (Gonzalez-Sandoval et al., 2015; Harr et al., 2016; Kind et al., 2013; Towbin et al., 2013; van Steensel and Belmont, 2017). Thus far, only these three H3K9me-based tethers, LBR, PRR14, and CEC-4, have been identified. Being a nonmembrane, nuclear lamina-associated protein, PRR14 is unique. However, the nature of PRR14 nuclear lamina association remained to be investigated.

Here, we have mapped a PRR14 nuclear lamina binding domain (LBD) that is both necessary and sufficient for nuclear lamina association, and also identified functional LBD core residues that are conserved beyond mammals. The discovery of a modular PRR14 LBD, in addition to the modular N-terminal HP1/heterochromatin binding domain, is consistent with the tethering function of PRR14. We also provide evidence that phosphorylation-dephosphorylation cycles within the LBD contribute to PRR14 dynamics at the nuclear lamina. Consistent with these results, we identified a function for the evolutionary conserved PRR14 C-terminal Tantalus in encoding a PP2A phosphatase recognition site (Hertz et al., 2016; Wang et al., 2016) that regulates PRR14 nuclear lamina association. The overall findings provide key insights into the mechanism and evolutionary conservation of the nuclear lamina attachment mechanism of the PRR14 tether.

## Results

### Identification of a minimal PRR14 domain that is sufficient for nuclear lamina association

We showed previously that the PRR14 N-terminal 1-135 region is necessary and sufficient for heterochromatin binding through a PRR14 <u>P</u>X<u>V</u>X<u>L</u> HP1/heterochromatin binding motif at positions 53-57 (LAVVL) (Fig. 1) (Poleshko et al., 2013). We also found that the C-terminal portion of PRR14 (residues 366-585) localized to the nucleus, but not to the nuclear lamina (Fig. 1C, D). When independently expressed, the highly conserved Tantalus domain localizes to the whole cell (Fig. 1C, D). To determine which region(s) of PRR14 are required, or sufficient, for nuclear lamina association, a series of C-terminal truncations of an N-terminal GFP-tagged 585 amino PRR14 protein were constructed (Figs. S1A). Nuclear lamina localization was found to be retained in HeLa cells for fragments that included the first 272 residues, but was lost with shorter fragments that included only the first 257 amino acids (Fig. S1A). With loss of nuclear lamina association, the residue 1-257, 1-241, 1-225 and 1-212 fragments appeared to localize to heterochromatin in perinucleolar regions and the peripheral heterochromatin compartment in HeLa cells (Fig. S1). To validate this interpretation, mutations were introduced in the LAVVL HP1/heterochromatin binding motif (LAVVLmut) in the 1-324 and 1-288 constructs (apparent nuclear lamina localization), and the 1-212 construct (apparent heterochromatin localization) (Fig. S1B). The LAVVLmut had no effect on the localization of the 1-324 and 1-288 fragments to the periphery, while the 1-212 fragment became nucleoplasmic as expected for loss of both nuclear lamina and heterochromatin binding (Fig. S1B). The localization to the nuclear lamina of the composite 1-324/LAVVLmut and 1-288/LAVVLmut proteins reinforces our previous interpretation that heterochromatin binding is not required for positioning of full length PRR14 at the nuclear lamina.

**Figure 1.**
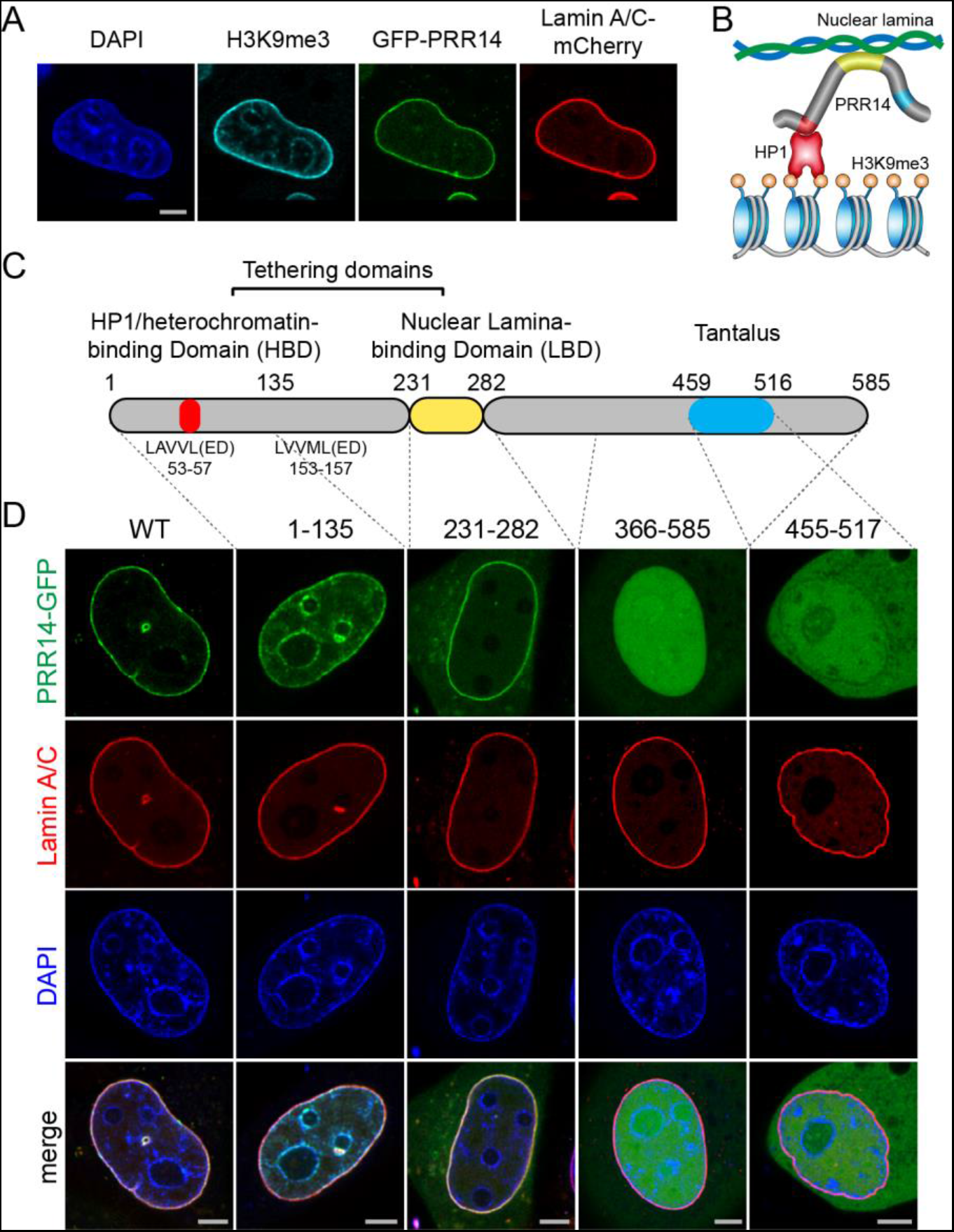
Identification of a modular PRR14 nuclear lamina binding domain. **(A)** Representative confocal imaging of HeLa cells showing localization of H3K9me3-marked heterochromatin (cyan), GFP-tagged PRR14 and mCherry-Lamin A/C (red). **(B)** A schematic model of H3K9me3/HP1 heterochromatin tethering to the nuclear lamina via PRR14. **(C)** A schematic representation of the PRR14 modular domain organization. **(D)** Representative confocal images of live HeLa cells transfected with GFP-tagged PRR14 fragments, as indicated. The N-terminal fragment (1-135) shows localization to heterochromatin, C-terminal fragment (366-585) localizes in nucleoplasm, centrally-located (231-282) fragment identified as a minimal nuclear lamina binding domain (LBD). Scale bars: 5μm.

Based on the retention of nuclear lamina association with the residue 1-272 fragment, a C-terminal border of a lamina binding domain (LBD) was provisionally assigned at position 282, providing a ten-amino acid buffer from the mapped endpoint. We found previously that the 1-135 region is an autonomous heterochromatin binding domain (Fig. 1C, D). As part of more comprehensive bioinformatic analysis of the PRR14 protein, we detected a degenerate duplication at the N-terminus corresponding to residues 9-100 and 106-214 (Fig. 1C). The duplicated region included a second candidate <u>P</u>X<u>V</u>X<u>L</u> motif at positions 153-157 (Fig. 1C). However, mutation of the LAVVL at positions 52-56 was sufficient to release the residue 1-212 fragment from heterochromatin (Fig. S1B). Although the second <u>P</u>X<u>V</u>X<u>L</u> motif does not function autonomously in heterochromatin binding, based on the duplication, the 1-212 region was viewed as relating to heterochromatin binding. Therefore, a 231-282 module was designed as a candidate LBD. PRR14 encodes an NLS near the N-terminus at positions 30-36 (Poleshko et al., 2013), and a surrogate SV40 NLS was therefore introduced at the N-terminus of the 231-282 fragment.

Strikingly, the 231-282 fragment localized at the nuclear periphery, tracing the nuclear lamina similarly to full length PRR14 (Fig. 1D). Localization of the 231-282 fragment to within the nucleus was confirmed using the benchmark differential permeability method, which in this case measures GFP-antibody nuclear/cytoplasmic accessibility of the GFP-LBD fusion protein (Fig. S2). As described below, we used evolutionary guidance and empirical fine mapping to identify additional contiguous downstream sequences between positions 283 and 351 that enhance nuclear lamina association of the 231-282 LBD.

### The PRR14 231-282 LBD is evolutionarily conserved beyond mammals

Initial alignments predicted that the PRR14 tether was unique to mammals (Poleshko et al., 2013). One difficulty in assessing the extent of evolutionary conservation of the PRR14 gene is the existence of the paralogous PRR14L gene (Table S1). The basis for assigning PRR14L (2,151aa versus 585aa for PRR14) as a human paralog is based on the highly conserved domain at the C-terminus of both proteins, positions 459-516 in PRR14 and positions 2026-2083 in PRR14L, which show 74% identity and 79% similarity in this region. As noted, this region has now been included as a protein family (Pfam) domain (El-Gebali et al., 2019), Tantalus, which was defined by homology with the Drosophila Tantalus protein (Fig. 1C) (Aravind and Iyer, 2012; Dietrich et al., 2001; El-Gebali et al., 2019). Based on overall lack of homology between mammalian PRR14, PRR14L, and Drosophila Tantalus, we currently believe that the Tantalus domain represents a shared module among these proteins with different functions. However, the presence of the Tantalus domain made it difficult to evaluate whether potential PRR14 orthologs in nonmammalian vertebrates were related to PRR14 or PRR14L (Table S1).

Having identified the human PRR14 231-282 LBD as a functional module, we focused our search for other proteins with similarity to the human LBD. We found evidence that the 231-282 LBD was present in PRR14-like proteins in reptiles and amphibians (Fig. 2A, B, Table S1). In view of these findings, a systematic evaluation of the functional conservation of the 231-282 LBD was initiated. As a first test, we examined mouse PRR14, where the LBD shares 94 percent sequence identity with the human PRR14 LBD (Fig. 2B). As expected, both the full length mouse PRR14 and human PRR14 proteins localized to the nuclear lamina in mouse cells (Fig. S3A). The gecko lizard and xenopus frog PRR14 proteins displayed 48 and 44 percent sequence identity and 59 and 55 percent sequence similarity within the human PRR14 231-282 LBD, respectively. The full length gecko and xenopus candidate PRR14 orthologs are larger than the 585 amino acid human PRR14 (727 and 699 amino acids, respectively), and show very limited homology overall with the human PRR14, outside of the Tantalus domain (82 and 70 percent identity, respectively) (Table S1). As further evidence of the functional conservation of the LBD, the full length human PRR14 was found to localize to the nuclear lamina in *Xenopus laevis* cells (Fig. S3A). To further test the modularity, and confirm functionalities of the LBDs, the human and xenopus LBDs were introduced into xenopus cells, and were found to localize to the nuclear lamina (Fig. S3B). Similarly, the LBD from the putative xenopus PRR14 protein localized to the nuclear lamina in human cells (Fig. S3B). As in Figure S2, we carried out differential antibody accessibility analysis to confirm that the xenopus GFP-tagged LBD localized inside the nucleus of HeLa cells (data not shown). The candidate gecko LBD was analyzed in detail below.

**Figure 2.**
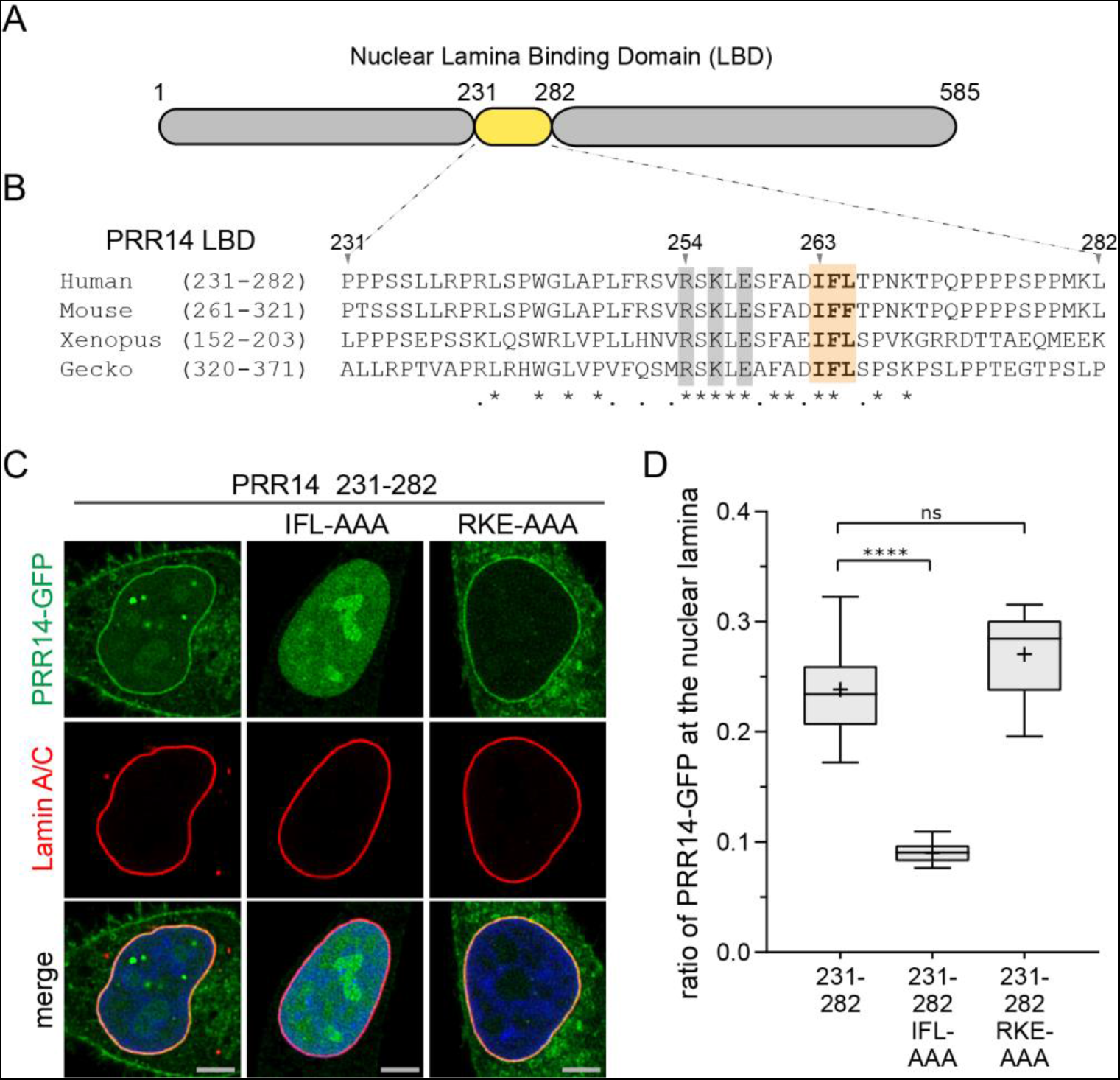
Identification of functional evolutionarily conserved residues in the 231-282 LBD. **(A)** PRR14 map depicting the minimal 231-282 LBD. **(B)** Protein sequence alignment of the human 231-282 PRR14 LBD with mouse, xenopus, and gecko identified conserved residues: charged (gray) and hydrophobic (orange). Amino acid sequence identity (*) and similarity (.) are indicted. **(C)** Representative confocal images of HeLa cells transfected with GFP-tagged PRR14 231-282 LBD containing the indicated amino acid substitutions. **(D)** Box plot demonstrating the proportion of indicated PRR14-GFP proteins at the nuclear lamina, calculated using Lamin A/C signal as a mask. Boxes: median, interquartiles range with Tukey whiskers (“+” mean value). n=20 cells per condition. Statistical analyses performed using one-way ANOVA test with Dunn’s multiple comparison; **** p<0.0001, ns: not significant. Scale bars: 5μm.

Having shown common functionality of nonmammalian LBDs, the alignment of the four LBD sequences was next used to predict key functional residues (Fig. 2A, B, S3C). A provisional core homology region in the human PRR14 LBD was identified at positions 254-265 (RSKLESFADIFL) (Fig. 2B, S3C, Table S1), and independent sets of alanine substitutions were introduced for the charged residues R, K, and E, and the hydrophobic residues I, F and L. Substitution of each of the RKE residues to A (RKE-AAA) had no effect on localization of the human PRR14 231-282 LBD, while substitution of the IFL residues with AAA resulted in dramatic loss of nuclear lamina localization of the LBD (Fig. 2C, D). These results were consistent with the loss of nuclear lamina localization observed for the residue 1-257 truncation, which retained the RKE constellation, but was missing the IFL motif (Fig. S1, S3C).

### The 231-282 LBD and conserved residues therein are required for efficient nuclear lamina localization of full length PRR14

The above results predicted that the 231-282 region would be necessary for localization of PRR14 to the nuclear lamina. HeLa cells were transfected with GFP-PRR14 carrying the deletion, del231-282. As shown in Figure 3, deletion of the 231-282 region resulted in significant loss of nuclear lamina localization. The IFL to AAA substitution was sufficient to produce a similar loss in nuclear lamina localization (Fig. 3B-D). The requirement for the IFL sequence was consistent with results using the modular 231-282 fragment (Fig. 2B-D). To assess whether the residual localization at the nuclear periphery was due to binding to peripheral heterochromatin, mutations were introduced in the LAVVL HP1/heterochromatin binding motif in the context of the del231-282 and IFL to AAA mutants. Heterochromatin binding was assessed in mouse cells which feature H3K9me3/HP1-rich chromocenters that decorate the nucleoli and periphery, as well as a peripheral layer of heterochromatin (Eberhart et al., 2013; Politz et al., 2013). As shown in Figure S4A, the composite LAVVLmut/del231-282 and LAVVLmut/IFL-AAA mutants lost chromocenter heterochromatin localization as expected, but retained residual peripheral localization. A similar experiment was carried out in HeLa cells, which showed that residual binding at the nuclear periphery could occur in regions barren of H3K9me3 heterochromatin (Fig. S4B). These results suggested that the 231-282 LBD is required for efficient nuclear lamina localization, but additional sequences were contributing nuclear lamina localization. However, extending the deletion on the N-terminal side (135-282) did not impact the residual nuclear lamina binding (data not shown).

**Figure 3.**
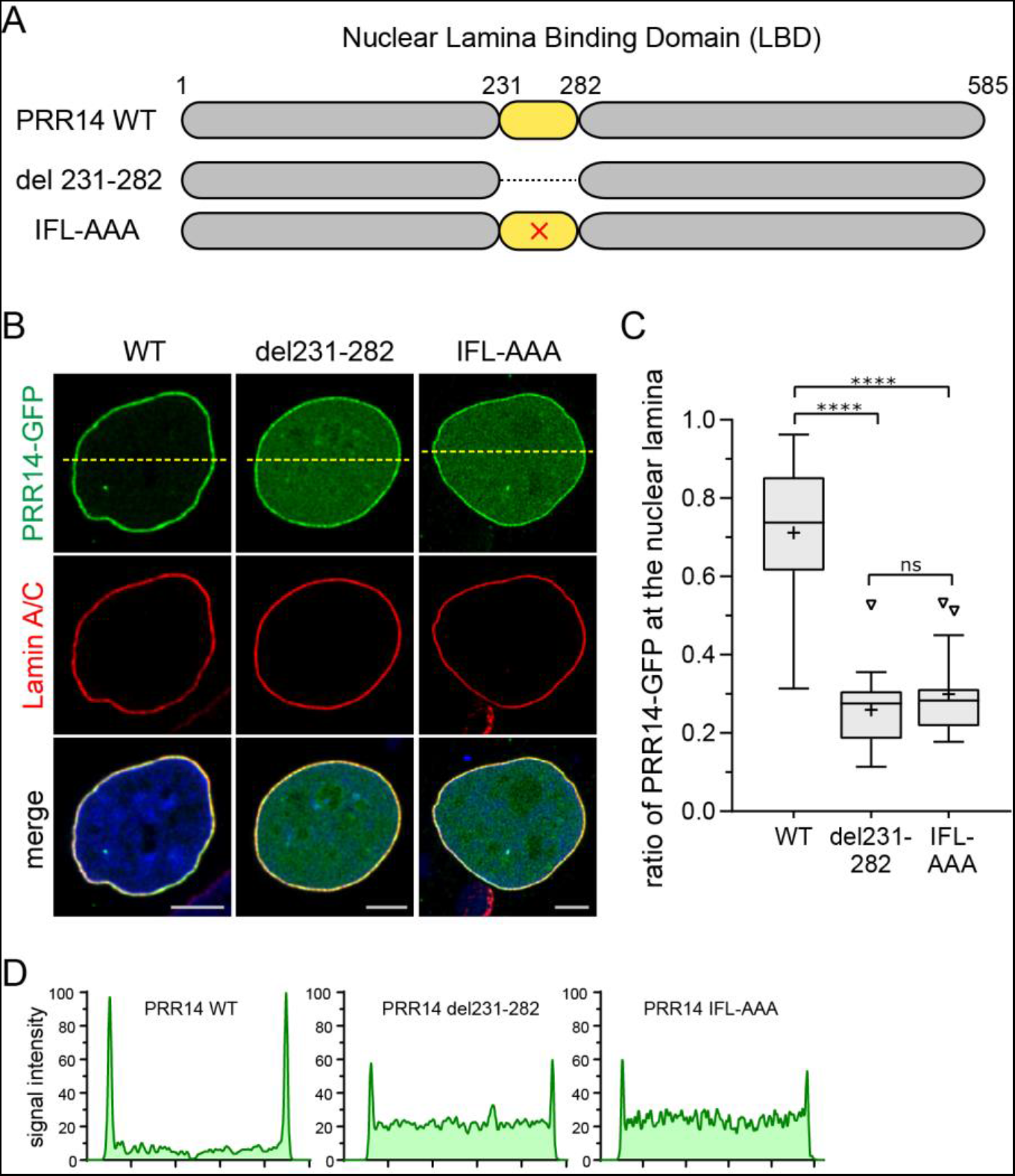
The 231-282 LBD is required for efficient localization of PRR14 to the nuclear lamina. **(A)** Map of human PRR14 depicting the 231-282 LBD, del231-282, and IFL to AAA substitution (x) in the conserved core sequence. **(B)** Representative confocal images of HeLa cells transfected with GFP-tagged constructs (green) depicted in Panel A, Lamin A/C (red), and counterstained with DAPI. **(C)** Box plot demonstrating the proportion of indicated PRR14-GFP proteins at the nuclear lamina, calculated using Lamin A/C signal as a mask. Boxes: median, interquartiles range with Tukey whiskers (“+” mean value). n=20 cells per condition. **(D)** Line signal intensity profiles of corresponding images in panel B indicated by dashed lines. Statistical analyses performed using one-way ANOVA test with Dunn’s multiple comparison; **** p<0.0001, ns: not significant. Scale bars: 5μm.

### Identification of an optimal, modular nuclear lamina binding domain

We hypothesized that residual nuclear lamina binding of the del231-282 mutant (Fig. 3B, D) could be through binding to partner proteins positioned at the nuclear lamina, or that the LBD might extend further to the C-terminal side. Alignment of human and gecko PRR14 sequences revealed several conserved downstream motifs (denoted B, C, D) that were candidates for contributing to nuclear lamina binding, along with motif A within the 213-282 LBD (Fig. 4A). Based on these findings, a human PRR14 231-351 fragment was designed that showed nuclear lamina association that was quantitatively indistinguishable from full length WT PRR14 (Fig. 4B, C). To determine the role(s) of motifs B, C, and D, fragments encompassing positions 231-324, 231-297, and 231-288 were designed. The 231-324 fragment again showed nuclear lamina association that was not significantly different from the full length protein, indicating that motifs B and/or C were important (Fig. 4B, C). As expected, the 231-288 fragment showed weaker association, similar to the 231-282 LBD. Surprisingly, the PRR14 283-351 fragment containing only motifs B and C localized to the nuclear lamina, although deletion of these sequences did not detectably reduce nuclear lamina binding in the initial mapping experiments (Fig. S1). To investigate the biological relevance of these motifs, three equivalent fragments from the putative gecko PRR14 protein were designed encompassing motifs A, A and B, or A, B, and C: 320-371, 320-410, and 320-444, respectively. Remarkably, all three gecko PRR14 fragments localized to the nuclear lamina in HeLa cells, with the 320-444 fragment being most efficient (Fig. S5). These results indicate that the mechanism of PRR14 nuclear lamina binding is conserved in vertebrate evolution and that motifs A, B, and C are required for optimal binding.

**Figure 4.**
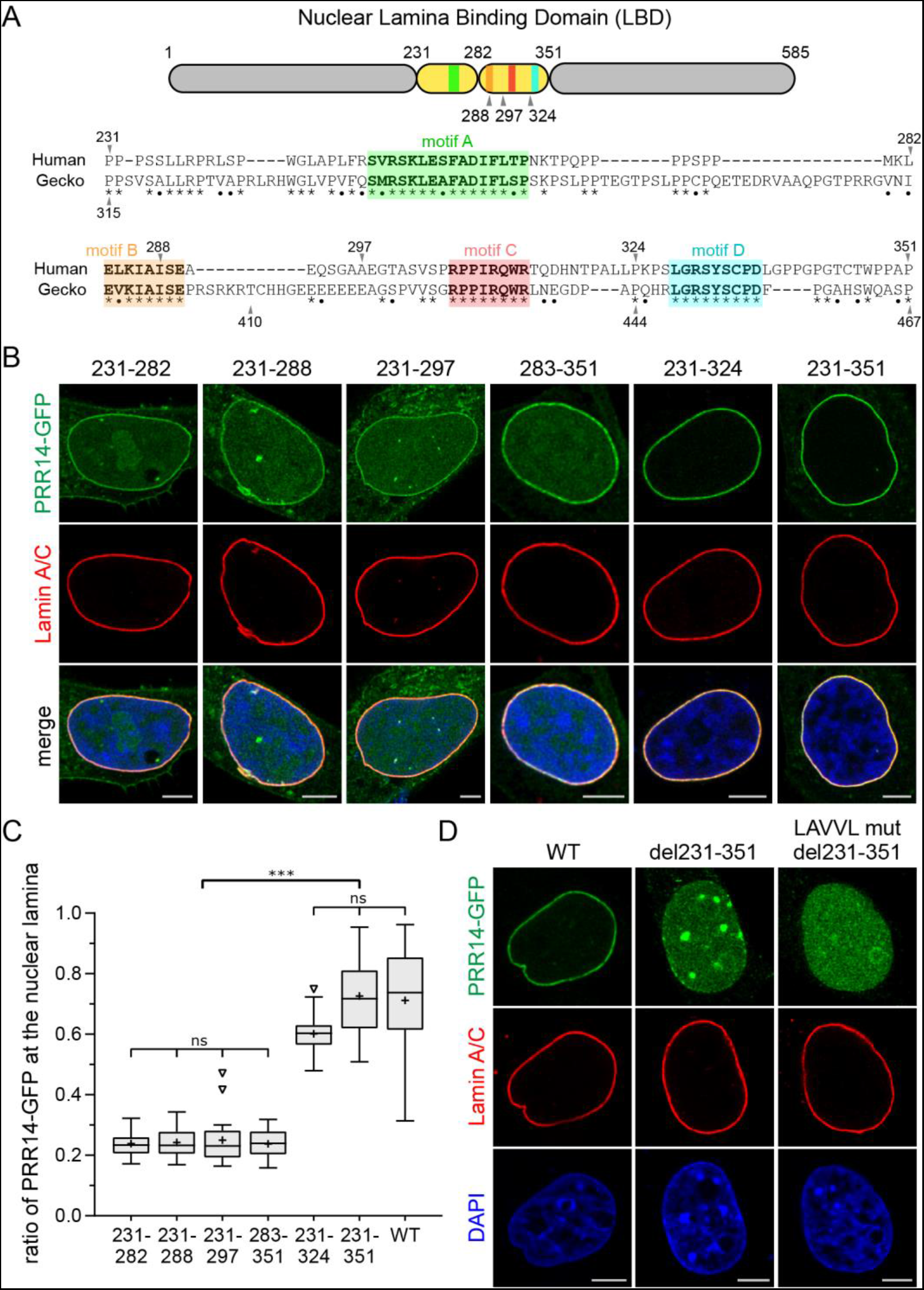
Identification of an optimal PRR14 LBD module. **(A)** Map, and sequence conservation of region downstream of 231-282 LBD hypothesized to contribute to nuclear lamina binding. Sequence alignment of human and gecko PRR14 identified candidate functional motifs. Amino acid sequence identity (*) and similarity (•) are indicted. Conserved motifs are designated as A through D. **(B)** Representative confocal images of HeLa cells transfected with indicated N-GFP tagged PRR14 fragments. Independent from the 231-282 LBD, the 283-351 fragment was found to localize to the nuclear lamina as a second independent modular LBD domain. **(C)** Box plot demonstrating the proportion of indicated PRR14-GFP proteins from panel B at the nuclear lamina, calculated using Lamin A/C signal as a mask. Boxes: median, interquartiles range with Tukey whiskers (“+” mean value). n=20 cells per condition. **(D)** Representative confocal images of HeLa cells transfected with indicated N-GFP tagged PRR14 constructs: WT, 231-351 deletion and composite mutant with 231-351 deletion and substitutions in the LAVVL sequence (LAVVLmut) required for heterochromatin binding. Statistical analyses performed using one-way ANOVA test with Dunn’s multiple comparison; *** p<0.001, ns: not significant. Scale bars: 5μm.

Having defined an extended modular LBD, we constructed a larger deletion in human PRR14 (del231-351) which was found to more completely disable nuclear lamina binding (Fig. 4D) compared to del231-282 (Fig. 3B, C and Fig. S4). Again, the del231-351 protein appeared to relocate to dense heterochromatin and this was confirmed by constructing a composite mutant LAVVLmut/del231-351 that localized throughout the nucleoplasm (Fig.4D). Based on analyses of isolated domains, we conclude that the 231-324 LBD is sufficient and optimal for nuclear lamina binding. These findings support the known tethering function of PRR14 by defining a novel modular domain capable of mediating bridging between the nuclear lamina and heterochromatin.

### Evidence for regulation of PRR14 nuclear lamina-association through LBD phosphorylation

Mitotic entry is accompanied by numerous phosphorylation events that trigger nuclear envelope and nuclear lamina disassembly (Dephoure et al., 2008). These phosphorylations are catalyzed by mitotic serine-threonine cyclin-dependent kinase (CDK), and the majority of phosphorylation sites conform to the consensus sequences serine-proline (SP) and threonine-proline (TP). We found previously that PRR14 is disassembled from the intact nuclear lamina during early mitosis (Poleshko et al., 2013). We therefore searched the PRR14 protein sequence for SP and TP sites, and also data-mined PhosphsitePlus (Hornbeck et al., 2015) and ProteomicsDB (Schmidt et al., 2017) data bases for evidence of in vivo SP/TP phosphorylation sites within the LBD that could regulate nuclear lamina association. In addition, we re-purposed our data from a BioID-based (Roux et al., 2012) search for PRR14 partners, were we could examine phosphorylation of the PRR14 bait protein (data not shown). As shown in Table S2, we identified four in vivo phosphorylation sites in the SP/TP context within the 231-282 LBD at positions S242, T266, T270 and S277. Of note, the S242 and S277 phosphorylation sites were detected in all data sets, and no additional sites were detected in the larger 231-324 LBD fragment.

We substituted all four (S/T) residues with a phosphomimetic residue, glutamic acid, in the context of the GFP-tagged 231-282 LBD fragment (Fig. 5A). This resulted in a dramatic loss of nuclear lamina localization, with accumulation in the nucleoplasm (Fig. 5B). Introduction of the four S/T glutamic acid phosphomimetic substitutions into the full length human PRR14 protein produced a similar loss of nuclear lamina association (Fig. 5D-F). Phosphomimetic substitutions are not always fully effective at reproducing the phosphorylated state (Dephoure et al., 2013), and therefore it is difficult to assess whether additional mechanisms account for the observed residual nuclear lamina retention.

**Figure 5.**
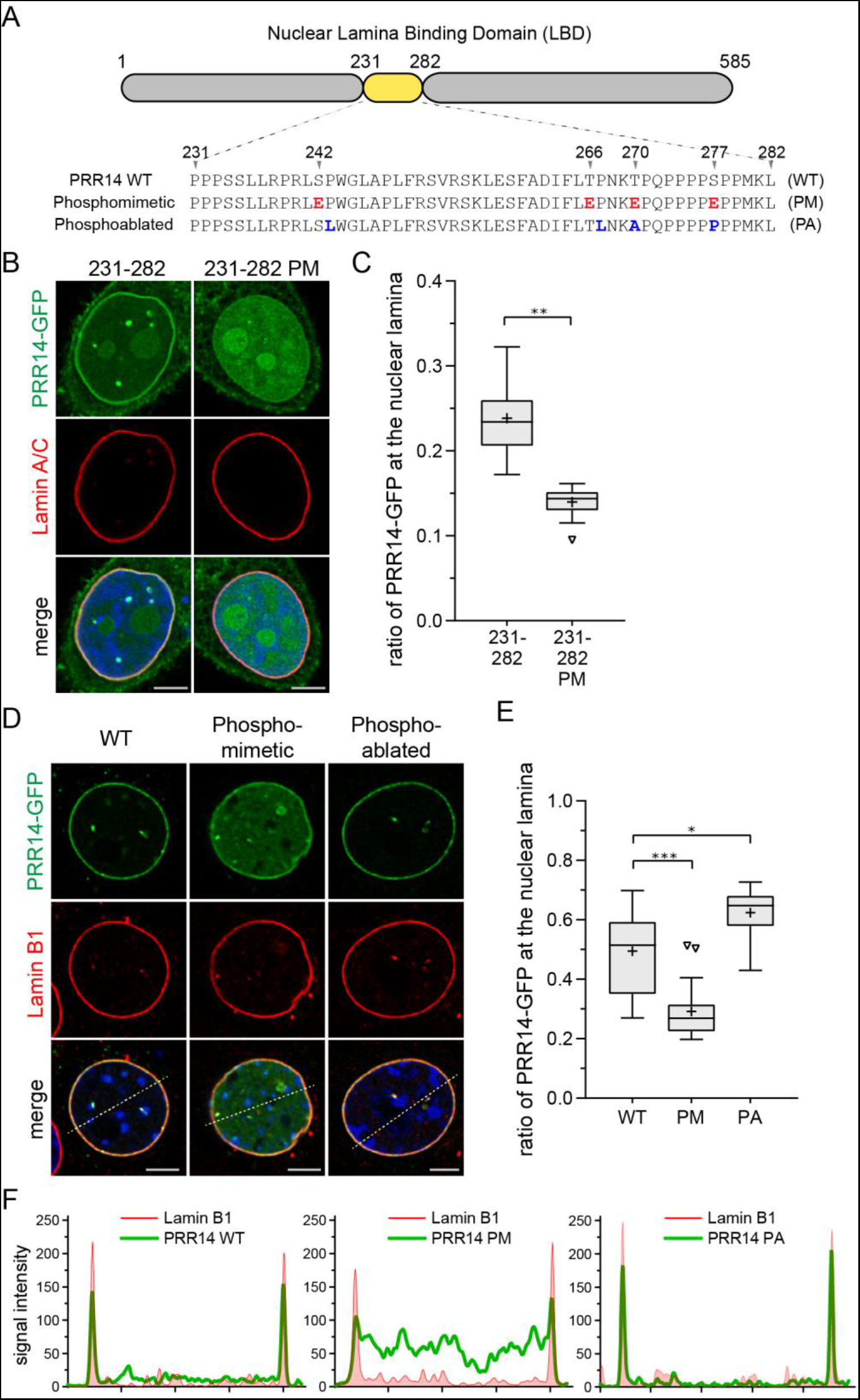
LBD phosphomimetic and phosphoablation substitutions affect PRR14 nuclear lamina association. **(A)** Map and sequence of the human PRR14 231-282 LBD highlighting in vivo serine/threonine CDK SP/TP phosphorylation sites at positions 242, 266, 270 and 277. Phosphomimetic (PM) and phosphoablation (PA) substitutions are shown. For phosphomimetic substitutions, the four S/T phosphorylation sites were collectively changed to glutamic acid. For phosphoablation, the SP/TP context at all four sites was disrupted using cancer-related mutations (Table 2) **(B)** Representative confocal images of HeLa cells express the GFP-tagged PRR14 LBD (231-282) phosphomimetic (PM) glutamic acid substitutions resulted in loss of nuclear lamina localization. **(C)** Box plot demonstrating the proportion of indicated PRR14-GFP proteins from panel B at the nuclear lamina, calculated using Lamin A/C signal as a mask. Boxes: median, interquartiles range with Tukey whiskers (“+” mean value). n=20 cells per condition. **(D)** Representative confocal images of C2C12 cells express phosphomimetic (PM) and phosphoablation (PA) GFP-tagged full length PRR14. PM substitutions resulted in loss of PRR14 nuclear lamina association, as seen with the 231-282 LBD. The PA substitutions appeared to result in stronger localization at the nuclear lamina. **(E)** Box plot demonstrating the proportion of indicated PRR14-GFP proteins from panel D at the nuclear lamina, calculated using Lamin B1 signal as a mask. Boxes: median, interquartiles range with Tukey whiskers (“+” mean value). n=20 cells per condition. **(F)** Line signal intensity profiles of corresponding images in panel D indicated by dashed lines. Statistical analyses performed using one-way ANOVA test with Dunn’s multiple comparison; *** p<0.001, ** p<0.01, * p<0.05, ns: not significant. Scale bars: 5μm.

Next, the four SP/TP sites were ablated in the context of the full length PRR14 protein (Fig. 5A). Data mining indicated that all four SP/TP sites were mutated in a variety of cancers (Table S2). Rather than ablating the four S/T residues with alanine residues, we introduced cancer-associated S/T and proline mutations that are predicted to disable the SP and TP sites (Table S2). The composite PRR14 mutations (P243L, T267L, T270A, S277P) resulted in a quantifiable, and unexpected phenotype, whereby PRR14 accumulated more intensely at the nuclear lamina during interphase (Fig. 5D-F).

### PRR14 dynamically associates with the nuclear lamina during interphase

The finding that ablating the PRR14 LBD phosphorylation sites resulted in apparent higher affinity for the nuclear lamina suggested that the PRR14 tether might dynamically associate with the nuclear lamina during interphase. To measure the mobility of PRR14 at the nuclear lamina, we use the conventional fluorescence recovery after photobleaching (FRAP) approach (Goldman et al., 2002). Cells were transfected with either GFP-PRR14 or GFP-Lamin A. Peripheral regions-of-interest were laser-bleached, and the recovery time was monitored. We confirmed that, as a component of the nuclear lamina framework, Lamin A was quite stable over the measured 5 minute period (Fig. 6, Video S1), as observed earlier (Goldman et al., 2002). In contrast, PRR14 was found to exchange rapidly at the nuclear lamina with a t1/2 of 6.4 seconds (Fig. 6, Video S2). Taken together with the phosphoablation-phosphomimetic results, one interpretation of these findings is that the rapid exchange of PRR14 at the nuclear lamina is phosphoregulated.

**Figure 6.**
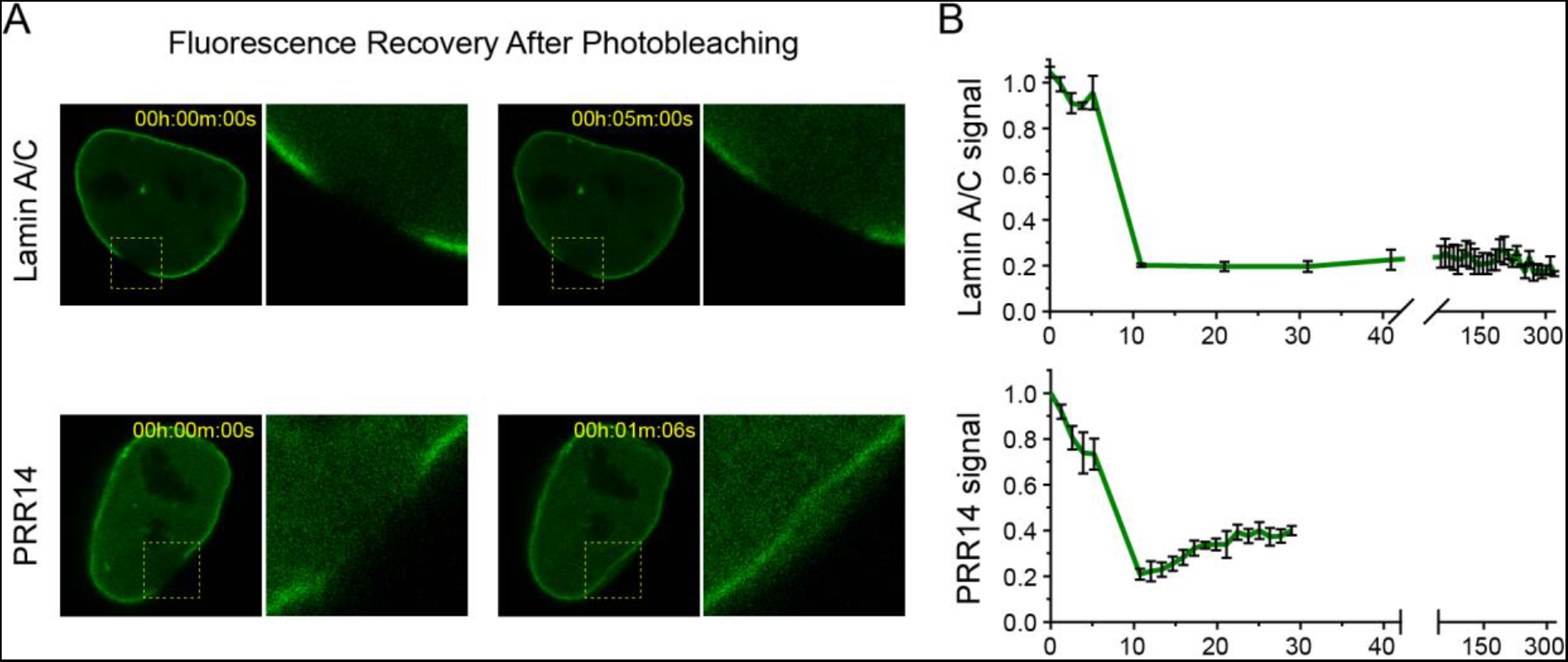
FRAP analysis showing that PRR14 is mobile at the nuclear lamina. HeLa cells were transfected with GFP-tagged Lamin A or GFP-tagged PRR14. A region of interest at the nuclear lamina was bleached and fluorescent recovery was monitored. **(A)** Representative confocal images of PRR14-GFP or LaminA/C-GFP after photobleaching and recovery. **(B)** Graph shows signal monitored over a 5 minute period for comparing PRR14 and Lamin A recovery times. X-axis shows time (s) prior to and after photobleaching. Lamin A was previously reported to have a recovery time of ca. one hour. PRR14 recovered completely within seconds, Data analyses using the EasyFrap software (Koulouras et al., 2018) indicated a t1/2 of 6.4 seconds with an R-square value of .95.

### PRR14 is recognized as a PP2A phosphatase substrate to regulate association with the nuclear lamina

After discovery of PRR14 in a functional screen, our data mining revealed that two interactor screens had detected PRR14, using HP1 (Nozawa et al., 2010; Rual et al., 2005) and phosphatase PP2A (Herzog et al., 2012) as bait proteins. PP2A is a complex trimeric phosphatase, alternatively composed of several isoforms of three subunit types: catalytic, structural and regulatory (Shi, 2009; Virshup and Shenolikar, 2009). This intricate organization allows recognition of a variety of substrates, thereby enabling PP2A to regulate a vast array of cellular functions.

As described above, PRR14 was recently identified as the human homolog of the drosophila Tantalus protein (Aravind and Iyer, 2012) and a PRR14 region near the C-terminus (459-516) was designated as the Protein Family (pfam) “Tantalus” domain (Fig. 7A). Data mining also revealed that the drosophila Tantalus protein, human PRR14, and the human paralog PRR14L, not only share the Tantalus domain, but all three proteins bind PP2A complexes (Glatter et al., 2009; Guruharsha et al., 2011; Herzog et al., 2012) (Fig. S6). Recently, a common motif was identified among PP2A substrates that is recognized by the PP2A B56alpha regulatory subunit (Hertz et al., 2016; Wang et al., 2016). We found that this motif (<u>L/F/M</u>xx<u>I</u>x<u>E</u>) (Hertz et al., 2016) corresponds to the most conserved portion of the Tantalus Pfam domain (Fig. 7, S6, Table S1). To test whether the PRR14 Tantalus domain could bind the PP2A B56alpha subunit, we performed a series of in vivo pulldown and mutagenesis experiments using a human PRR14 Tantalus domain fragment (455-517) and the human B56alpha subunit (Fig. 7B-E). Substitution of predicted key residues (Hertz et al., 2016) of the PRR14 Tantalus motif <u>F</u>ET<u>I</u>F<u>E</u> with alanine, resulted in a loss of binding to B56alpha (Fig. 7B-D), while substitution of two nonconserved residues (N483S, K484R) had no effect (Fig. 7C). Also, substitutions of the IY residues in Tantalus upstream conserved region (IYT) (Fig. S6) had no effect on B56alpha binding (data not shown), suggesting that these conserved residues have a distinct function. Interestingly, an F to L mutation in the first position of the PRR14 Tantalus consensus resulted in tighter binding to B56alpha, as predicted (Fig. 7D) (Hertz et al., 2016) (see Discussion). Reciprocal mutations in the B56alpha heat repeat binding pocket that engages the <u>L/F/M</u>xx<u>I</u>x<u>E</u> motif (Hertz et al., 2016) also resulted in loss of Tantalus binding (Fig. 7E).

**Figure 7.**
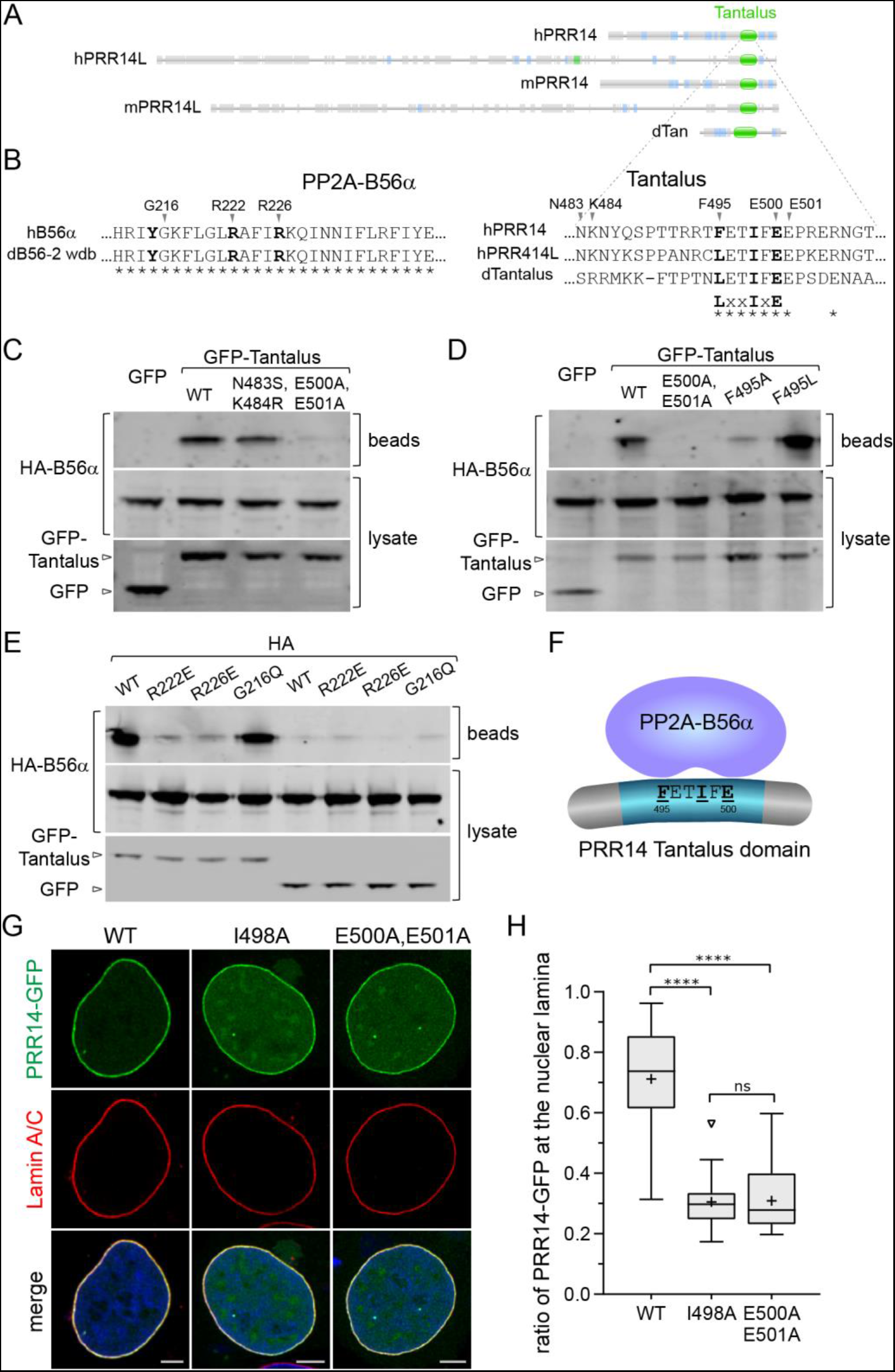
Pulldown, mutagenesis, and imaging experiments demonstrate that human PRR14 nuclear lamina localization is regulated by PP2A. **(A)** Maps extracted from Pfam (El-Gebali et al., 2019) comparing human and mouse PRR14 and PRR14L paralogs, and the Drosophila Tantalus protein, showing the relationship with the Tantalus domain. **(B)** Summary of mutations to test the role of the putative conserved PP2A B56alpha recognition motif L/F/MxxIxE in the Tantalus domain of human PRR14. On the left are shown residues (in bold) in the B56alpha heat repeat that are known to mediate recognition of the substrates encoding the L/F/MxxIxE motif (Hertz et al., 2016). The Drosophila B56alpha ortholog (dB56-2) shows 100 percent identity with human B56alpha in this region, consistent with the detection of Drosophila Tantalus-Drosophila B56alpha interactions (Guruharsha et al., 2011) (Fig. S6). On the right are shown residues (in bold) in the PRR14 Tantalus domain corresponding to the L/F/MxxIxE B56alpha recognition motif. Shown are positions mutated in B56alpha and PRR14 to test effects on interactions. Amino acid sequence identity (*) is indicted. **(C)** HeLa cells were cotransfected with human HA-tagged B56alpha and non-fused GFP, or GFP fused to the human PRR14 Tantalus domain. The GFP bait proteins (GFP only, GFP-Tantalus) were collected using anti-GFP beads and analyzed by western for interaction with HA-tagged B56alpha. GFP bait proteins were detected using anti-GFP antibodies and prey proteins were detected with anti-HA antibodies. As shown, GFP-Tantalus pulled down HA-B56alpha, while the control non-fused GFP did not. Amino acid substitutions of non-conserved PRR14 Tantalus residues N483 and K484 (see Panel B) had no effect, while substitution of E500 and E501 with alanine eliminated interaction as predicted (Hertz et al., 2016). **(D)** Experimental design was as in Panel C. Substitution of PRR14 Tantalus F495 with alanine inhibited binding, while substitution with leucine increased binding as predicted (Hertz et al., 2016). **(E)** Experimental design was as in Panel C. Amino acid substitutions were introduced into the HA-tagged B56alpha subunit that were predicted to have no effect (G216Q) or inhibit binding (R222E and R226E) to the GFP-Tantalus domain (Hertz et al., 2016). Results were consistent with these predictions. **(F)** A schematic cartoon showing PP2A interaction with PRR14 Tantalus domain. **(G)** Representative confocal images of HeLa cells transfected with indicated Tantalus domain amino acid substitutions introduced into full length GFP-tagged human PRR14 (see panel B). **(H)** Box plot demonstrating the proportion of indicated PRR14-GFP proteins from panel F at the nuclear lamina, calculated using Lamin A/C signal as a mask. Boxes: median, interquartiles range with Tukey whiskers (“+” mean value). n=20 cells per condition. Statistical analyses performed using one-way ANOVA test with Dunn’s multiple comparison; **** p<0.0001, ns: not significant. Scale bars: 5μm.

We hypothesized that PP2A regulates PRR14 localization at the nuclear lamina by dephosphorylating S/T residues in the LBD. It was therefore predicted that disabling the PRR14 <u>F</u>ET<u>I</u>F<u>E</u> motif would result in loss of PP2A binding to PRR14, an increase in LBD phosphorylation of the full length PRR14, and reduction nuclear lamina association. As shown Figure 7G-H, this was indeed the case, as tested with two independent mutations (I498A and E500A,E501A). Kinase and PP2A phosphatase activities are thereby implicated in mediating dynamic association of PRR14 with the nuclear lamina during interphase.

## Discussion

There is considerable interest in understanding how the heterochromatin compartment is organized at the nuclear envelope, and how lineage-inappropriate genes are silenced through such positioning (Amendola and van Steensel, 2014; Buchwalter et al., 2019; Gordon et al., 2015; Gruenbaum and Foisner, 2015; Harr et al., 2016; Lemaitre and Bickmore, 2015; Meister and Taddei, 2013; Padeken and Heun, 2014; Poleshko et al., 2017; Politz et al., 2013; Shevelyov and Nurminsky, 2012; Shevelyov and Ulianov, 2019; Ungricht and Kutay, 2015; van Steensel and Belmont, 2017; Wong and Reddy, 2015; Yanez-Cuna and van Steensel, 2017; Zullo et al., 2012). Emerging research findings have pointed to an evolutionarily conserved mechanism whereby a class of three “tethering proteins” function to organize H3K9me-modified heterochromatin at the nuclear periphery (Gonzalez-Sandoval et al., 2015; Harr et al., 2016; Kind et al., 2013; Poleshko and Katz, 2014; Poleshko et al., 2013; Towbin et al., 2013; van Steensel and Belmont, 2017). A common element of this class of tethers is that H3K9me heterochromatin modifications serve as anchoring points for attachment of heterochromatin to the nuclear periphery. Two members of this class, LBR (Olins et al., 2010) and CEC-4 (Gonzalez-Sandoval et al., 2015; Harr et al., 2016; Shevelyov and Ulianov, 2019) anchor heterochromatin to nuclear membranes, and their roles in heterochromatin organization, as well as cell differentiation have been described (Gonzalez-Sandoval et al., 2015; Solovei et al., 2013).

We identified human PRR14 protein as an epigenetic repressor and determined that it functions as a heterochromatin tether through HP1/H3K9me3, similarly to LBR (Poleshko and Katz, 2014; Poleshko et al., 2014; Poleshko et al., 2013). However, unlike the membrane-associated LBR and CEC-4 proteins, PRR14 is a nonmembrane protein and associates with the nuclear lamina through Lamin A/C (Poleshko et al., 2013). PRR14 is thereby expected to be more mobile than the membrane bound LBR (Giannios et al., 2017) and CEC-4 proteins. In this sense, PRR14 can be viewed as a heterochromatin-binding protein that associates with the nuclear lamina. The questions of: i) how PRR14 associates with the nuclear lamina, ii) the evolutionary conservation of the mechanism, and iii) how this association may be regulated, are the subjects of the work presented here.

Strikingly, we initially identified a minimal autonomous PRR14 LBD that mapped between positions 231-282 that was sufficient to target GFP to the nuclear lamina (Fig. 1C, D). The existence of PRR14 modular heterochromatin (1-135) (Poleshko et al., 2013) and nuclear lamina binding domains is consistent with the role of PRR14 as a tether (Fig. 8). Furthermore, these findings represent the first example of a modular mechanism through which HP1/H3K9me3 heterochromatin can be tethered to the nuclear lamina, rather than the inner nuclear membrane.

**Figure 8.**
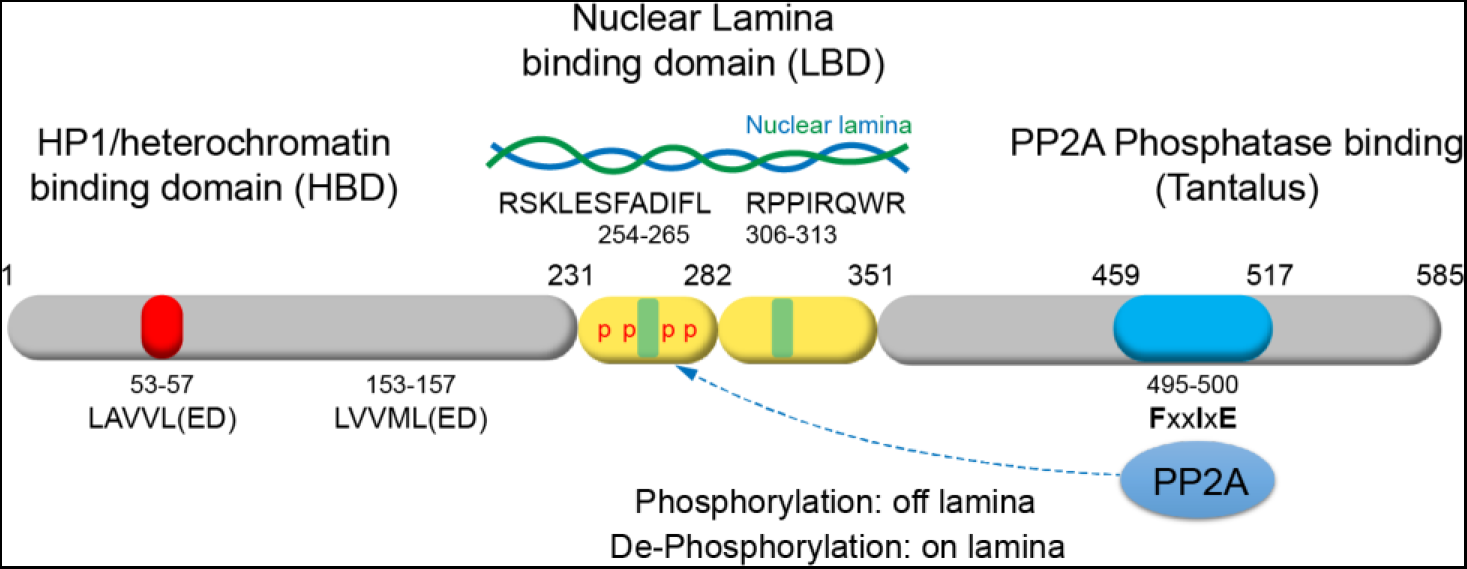
A schematic model of modular organization of the PRR14 functional domains. PRR14 is found to be highly modular protein. Several short evolutionary conserved motifs play key roles in bivalent tethering between heterochromatin and the nuclear lamina, and involved in regulation of association with the nuclear lamina.

Having identified a functional LBD, we were able to discriminate true PRR14 orthologs, versus PRR14L orthologs that do not show homology to the LBD (Table S1). Alignment of the human, xenopus, and gecko LBDs revealed a provisional core of conserved residues: RSKL(A/S)FA(D/E)IFL (Fig. 2B, Table S1). Mutation of conserved residues (IFL) in the human 231-282 LBD motif resulted in loss of nuclear lamina binding in the context of both the LBD domain and full length PRR14 (Figs. 2 and 3). Reciprocal interspecies nuclear lamina localization of full length PRR14 and the modular LBDs (Figs. S3, 5) suggests an evolutionary conserved attachment mechanism. Additional deletion and domain localization studies identified an optimal human PRR14 231-324 LBD that contained additional conserved motifs (Fig. 4). The relevance of these motifs was confirmed as the gecko 315-444 LBD showed optimal nuclear lamina localization among the fragments tested (Fig. S5). Despite a large body of evidence that PRR14 associates with the nuclear lamina, we have not yet determined whether PRR14 is among the proteins that interact directly with Lamin A/C (Simon and Wilson, 2013), the candidate partner identified by siRNA knockdown (Poleshko et al., 2013). However, the fact that human PRR14 interacts with the nuclear lamina in xenopus cells suggests that the fundamental mechanism for PRR14 nuclear lamina association is highly conserved. This fits well with the conservation of Lamin A between humans and xenopus, 73 percent identity and 87 percent similarity.

A general paradigm is that phosphorylation promotes disassembly of nuclear envelope components during mitosis, and that dephosphorylation is required for nuclear reassembly (Dephoure et al., 2008). Data mining, and our own mass spec analysis, identified four SP/TP phosphorylation sites within the 231-282 LBD that were candidates for regulating PRR14 nuclear lamina association (Table S2). Phosphomimetic mutations of these four SP/TP sites resulted in decreased association of both full length PRR14 and the 231-282 domain with the nuclear lamina (Fig. 5). Surprisingly, we found that ablating the four ST/TP sites resulted in a constitutive increase in association of PRR14 with the nuclear lamina during interphase (Fig. 4). One interpretation of these results is that pervasive phosphoregulation of PRR14-nuclear lamina association is manifested during interphase, as mediated by phosphorylation-dephosphorylation cycles. To test directly whether PRR14 dynamically associates with the nuclear lamina during interphase, we performed FRAP analysis, and found that PRR14 was highly mobile in comparison to the nuclear lamina component Lamin A (Fig. 6). We note that the LBD phosphomimetic substitutions promoted nucleoplasmic accumulation of PRR14, suggesting that phosphorylation drives a nucleoplasmic form. Taken together, the FRAP and mutagenesis experiments suggests that PRR14 dynamically reassociates between the nuclear lamina and nucleoplasm during interphase. We further speculate that the interphase-acting CDK2 SP/TP kinase (Hochegger et al., 2008) might contribute to these interphase dynamics. Future studies will be required to obtain more direct evidence for phosphoregulation of PRR14-nuclear lamina interactions.

Independent experiments, however, provided additional evidence for phosphoregulation of PRR14 association with the nuclear lamina. The conservation of a motif in the PRR14 Tantalus domain (Aravind and Iyer, 2012), and several phosphatase PP2A-binding partner studies (Glatter et al., 2009; Guruharsha et al., 2011; Herzog et al., 2012), pointed to a functional link between the Tantalus domain and PP2A (Figs. 7A, Fig. S6). For example, Drosophila PP2A was found to interact with the Drosophila Tantalus protein (Guruharsha et al., 2011), and the most conserved motif in the Tantalus domain was subsequently determined to specify PP2A substrates through the B56alpha subunit (<u>L/F/M</u>xx<u>I</u>x<u>E</u>) (Hertz et al., 2016). This short linear interacting motif (SLIM) is present in PRR14, PRR14L, and Drosophila Tantalus (Table S1, Fig. 6). Notably, the recent study by Hertz et al. detected PRR14 as a partner of B56alpha. We found that the PRR14 Tantalus domain was sufficient to bind the B56alpha subunit, and this interaction was sensitive to predicted amino acid substitutions in both the Tantalus domain and B56alpha (Fig. 7) (Hertz et al., 2016). Interestingly, comparison of the mammalian PRR14 and PRR14L Tantalus domains revealed an <u>F</u>xx<u>I</u>x<u>E</u> motif for PRR14 orthologs and an LxxIxE motif for PRR14L orthologs (Table S1, Fig. S6). As reported (Hertz et al., 2016) we found that an F to L substitution in the PRR14 motif enhanced binding to B56alpha (Fig. 7). These results fit well with the proposal that variations in this in the motif can modulate affinity for B56alpha (Hertz et al., 2016). Importantly, disabling of the PRR14 <u>F</u>xx<u>I</u>x<u>E</u> motif resulted in reduced PRR14 nuclear lamina association, consistent with loss of PP2A binding, and concomitant hyperphosphorylation of the LBD. These results support a role for PP2A regulating PRR14 association with the nuclear lamina during interphase (Fig. 8).

Previously, we found that PRR14 knockdown caused nuclear shape defects, while the dynamic behavior of PRR14 detected here suggests that PRR14 is not an intrinsic structural component of the nuclear lamina. How else might PRR14 contribute to nuclear structure? Perhaps PRR14 may provide a bridge through which the nuclear lamina can properly assemble on a base layer of heterochromatin at the end of mitosis. This function fits well with the idea that organization of heterochromatin at the nuclear periphery may contribute to nuclear structure (Bustin and Misteli, 2016; Stephens et al., 2019). But what then might be the functional relevance of the observed dynamic behavior of PRR14 at the nuclear lamina during interphase? An important breakthrough relevant to the peripheral heterochromatin was the mapping of genomic DNA sequences in close contact with the nuclear lamina (Guelen et al., 2008; Peric-Hupkes et al., 2010). These sequences, denoted lamina associated domains (LADs) have been found to constitute 35%-40% of nuclear DNA. It appears that LADs are contained in the peripheral heterochromatin compartment that can be visualized microscopically. Single cell studies have detected dynamic changes in LADs (Kind et al., 2013; Yanez-Cuna and van Steensel, 2017). It is possible that a dynamic tether may facilitate observed exchanges between the peripheral and perinucleolar heterochromatin compartments (Kind et al., 2013; van Koningsbruggen et al., 2010; van Steensel and Belmont, 2017; Vertii et al., 2019). The peripheral-perinucleolar exchanges are presumed to take place through mitosis, but a dynamic tether could facilitate more rapid exchanges during interphase. Alternatively, PRR14 mobility may simply reflect its association with the droplet-forming HP1 protein (Strom et al., 2017; Tatarakis et al., 2017).

As described, PRR14L is a much larger protein than PRR14, and does not encode identifiable tether domains, that is a <u>P</u>X<u>V</u>X<u>L</u> HP1-binding domain and a LBD (Table S1, Fig.7A). As such, PRR14L is not predicted to function as a tether. Rather, PRR14L seems to share largely the Tantalus domain and B56alpha binding motif with PRR14. As noted, the PRR14L motif (LxxIxE) is predicted to have higher affinity for B56alpha. Recent studies (Chase et al., 2019) identified PRR14L as a disease gene, driving age-related clonal hematopoiesis and contributing to myeloid neoplasia. The authors determined that human chromosome 22-acquired uniparental disomy (aUPD) created biallelic C-terminal truncations of PRR14L. As we have presented here in detail (Table 1), the authors similarly concluded that the paralogous relationship between PRR14 and PRR14L is defined solely by the common C-terminal Tantalus domain. Furthermore, they show that PRR14L accumulates at midbodies and suggest that it may play a role in cell division. We further suggest that PRR14L may function as a large scaffold and position PP2A through the Tantalus domain.

Our findings show that the 52 amino acid 231-282 LBD is sufficient to target the GFP protein to the nuclear lamina, while the 94 amino acid 231-324 fragment localizes as strongly as the full length protein. The modular LBD may therefore be useful for continued proof-of-concept experiments to test whether positioning of genes at the nuclear lamina is sufficient for gene silencing. If this paradigm holds, it may be possible to program gene silencing by conditional delivery of genes to the nuclear periphery. One obvious approach would be to use enzymatically disabled CAS9 fused with the LBD to deliver genes to the periphery using locus-specific guide RNAs.

In summary, PRR14 was found previously to function as a heterochromatin-nuclear lamina tether and here we show that PRR14 encodes a modular, evolutionarily conserved LBD. These findings reinforce the tethering model by demonstrating a mechanism by which PRR14 localizes to the nuclear lamina. We also provide evidence that the PRR14 tether exchanges rapidly at the nuclear lamina through phosphoregulation of the LBD, and that phosphatase PP2A plays a role in this process. Further study of tethering proteins will likely contribute to an understanding of the mechanisms that underlie heterochromatin disorganization in cancer and aging.

## Materials and methods

### Cells

Murine C2C12 skeletal myoblast and HeLa cells were received from the American Type Culture Collection (ATCC, cat# CRL-1772 and cat# CCL-2) and were tested negative for mycoplasma contamination. C2C12 and HeLa cells were maintained at 37°C in DMEM supplemented with 10% FetalPlex serum complex (Gemini, cat# 100-602), penicillin, and streptomycin. Alternatively, HeLa cells were grown in DMEM 10% FBS, supplemented with Penicillin-Streptomycin (Corning, cat# 30-002-CI), and Fungizone (ThermoFisher, cat# 15290-018). *Xenopus laevis* S3 cells were a gift from Matthew Good, University of Pennsylvania School of Medicine and were grown in L-15 Medium supplemented with 10% FBS, Sodium Pyruvate, and Penicillin-Streptomycin at 27°C.

### Plasmids

A human PRR14 expression vector was obtained previously from OriGene Technologies, Inc., encoding a C-terminal fusion with TurboGFP (cat# RG208696). The human PRR14 orf was transferred to N-terminal mGFP and mRFP vectors from OriGene (pCMV6-AN-mGFP, cat# PS100048, pCMV6-AN-mRFP, cat# PS100049) via the Origene PrecisionShuttle system using SgfI and MluI restriction sites. The N-terminal full length mGFP PRR14 fusion was used in this paper as a base construct for mutagenesis. The mouse PRR14 expression vector was obtained from Origene Technologies, Inc., encoding a C-terminal fusion with TurboGFP (cat# MG209414). The mouse PRR14 orf was transferred to the N-terminal mGFP vector from OriGene (pCMV6-AN-mGFP, cat# PS100048) using the Origene PrecisionShuttle system. For analyses of candidate modular LBDs from human, xenopus, and gecko PRR14, and for expression of the isolated Tantalus domain, gene synthesis was used (Genewiz). LBDs and Tantalus sequences were synthesized with terminal SgfI and MluI restriction sites to facilitate cloning into the OriGene pCMV6-AN-mGFP vector using the PrecisionShuttle system. The candidate LBD modules were designed in most cases to include an SV40 NLS (PKKKRKV) at the N-terminal side to enhance nuclear import, with the final configuration being mGFP-NLS-LBD. The empty OriGene pCMV6-AN-mGFP vector was used as a GFP-only control, as needed. As a reference for the nuclear lamina, mCherry-LaminA-C-18 was used, which was a gift from Michael Davidson (Addgene plasmid # 55068; http://n2t.net/addgene:55068; RRID:Addgene_55068). For FRAP experiments pEGFP-D50 lamin A was used, which was a gift from Tom Misteli (Addgene plasmid # 17653; http://n2t.net/addgene:17653; RRID:Addgene_17653) (Scaffidi and Misteli, 2005). For Tantalus-PP2A B56alpha pulldown experiments, an HA-tagged human B56alpha expression vector was used, a gift from David Virshup (Addgene plasmid # 14532; http://n2t.net/addgene:14532; RRID:Addgene_14532) (Seeling et al., 1999).

### Site-directed mutagenesis

Site-directed mutagenesis was carried out using the Agilent Technologies QuikChange II XL Site-Directed Mutagenesis Kit (cat# 200521). Mutagenic primers were designed using the Agilent web-based QuikChange Primer Design Program (www.agilent.com/genomics/qcpd). Mutagenic primers were purchased from Integrated DNA Technologies, Inc. (IDT). Mutagenic primer sequences are provided:

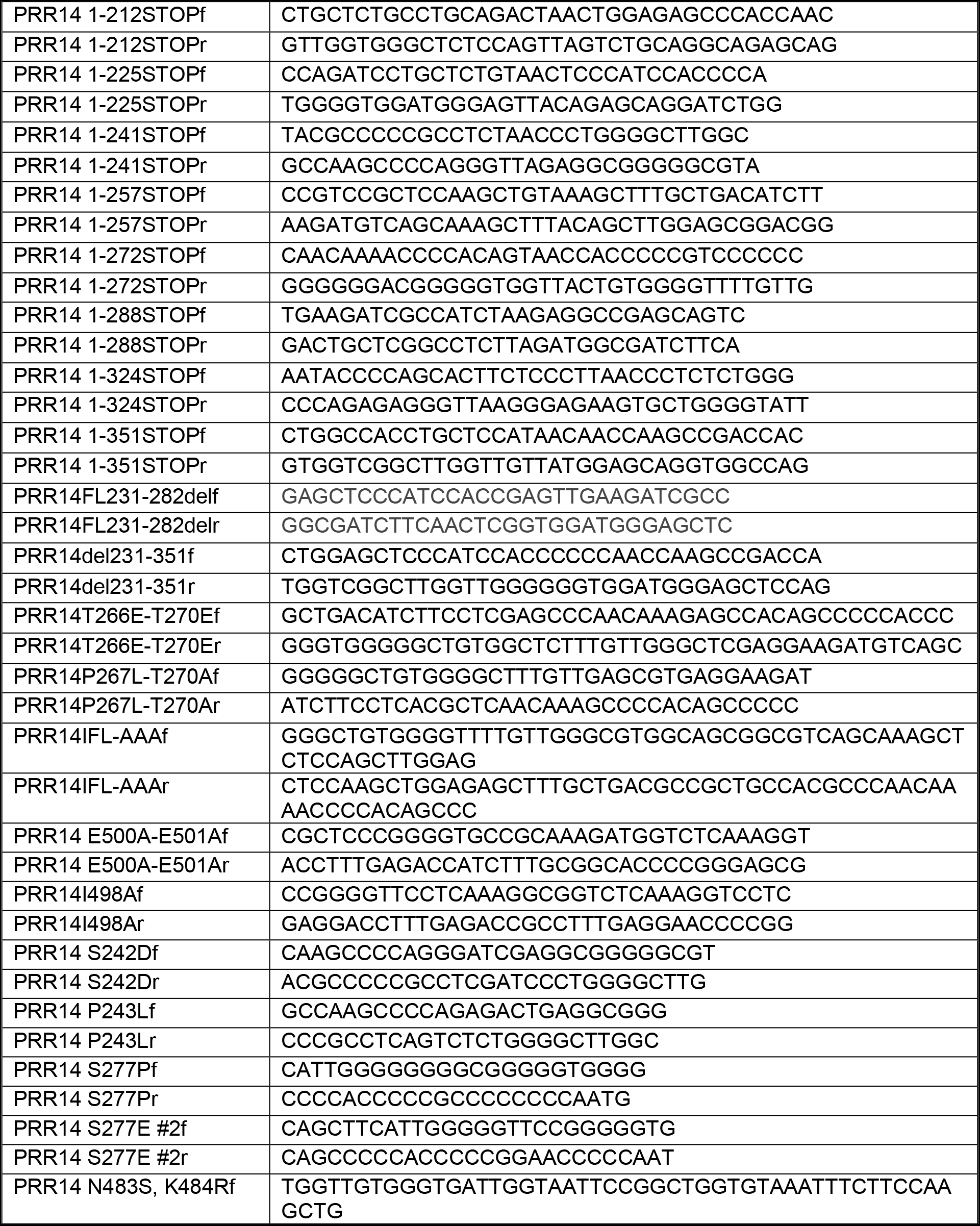
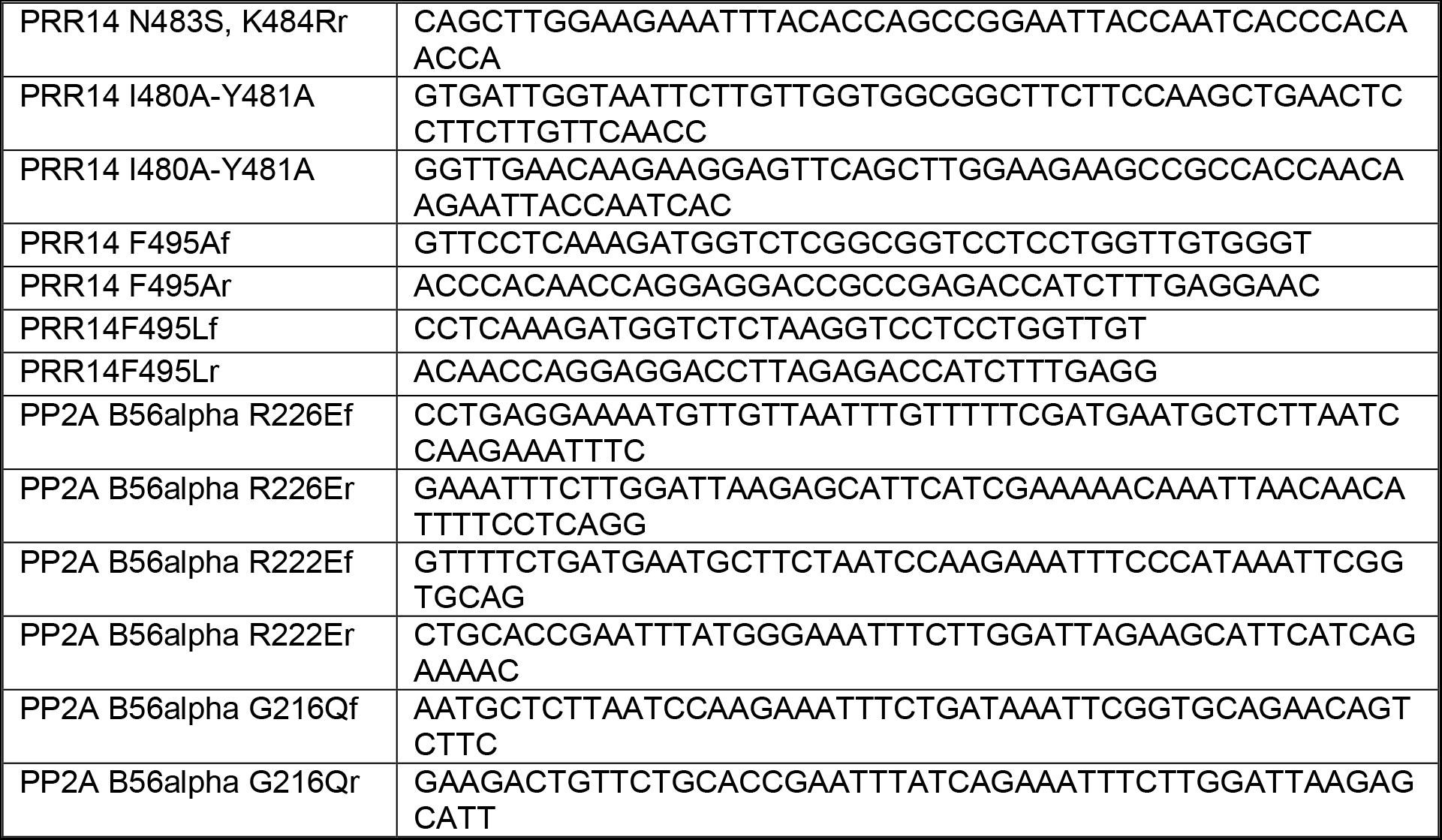

### Transfection

Plasmid transfections were performed using Lipofectamine 3000 (Invitrogen, cat# L3000008) according to manufacturer’s instructions. For some experiments, Lipofectamine 2000 was used (Invitrogen, cat# 11668030) according to the manufacturer’s instructions. To create HeLa cells stably expressing PRR14-mRFP or Lamin A/C-mCherry, HeLa cells were transfected with pCMV6-AN-mRFP-PRR14 or mCherry-LaminA-C-18, respectively using Lipofectamine 2000 according to the supplier’s protocol, and then were selected with G418 (ThermoFisher, cat# 10131035) 48 hours post transfection. Next cells were FACS sorted to enrich for mRFP/mCherry positive cells. For confocal imaging cells were plated on 8-well idibi µ-slides (Ibidi, cat# 80826) or glass bottom 35mm culture dish (MatTek, cat# P35G-1.0-14-C), then transfected at 50% confluency and fixed 24 h post-transfection or imaged live. Xenopus S3 cells were transfected in 35mm MatTek glass bottom dishes. Cells were transfected at 60-80% confluency using Fugene HD reagent (Roche, cat# E2311). Fugene HD (3 μl) was added to 400 μl Opti-MEM media, along with 1 μg DNA, the mix was incubated for 20 min. Next 400 μl Opti-MEM was added and the 800 μl volume was applied to the dish. Cells were cultured at 27°C and imaged 48 h post-transfection.

### Immunofluorescence

HeLa and C2C12 cells were fixed with 2% paraformaldehyde (PFA) (EMS, cat# 15710) for 10 min at RT, washed 3 times with DPBS (Gibco, cat#14190-136), then permeabilized with 0.25% Triton X-100 (ThermoFisher, cat# 28314) for 10 min. After permeabilization, cells were washed 3 times with DPBS for 5 min, then blocked in 1% BSA (Sigma, cat# A4503) in PBST (DPBS with 0.05% Tween 20, pH 7.4 (ThermoFisher, cat# 28320) for 30-60 min at RT. Next, samples were incubated with primary antibodies for 1 hour at RT, washed three times with PBST for 5 min, and then incubated with secondary antibodies for 30-60 min at RT followed by washing two times with PBST for 5 min. Samples were counterstained with DAPI solution (Sigma-Aldrich, cat# D9542) for 10 min at RT, then rinsed with PBS. Slides were mounted using 80% glycerol mounting media: 80% glycerol (invitrogen, cat#15514-011), 0.1% sodium azide (Sigma-Aldrich, cat# S2002), 0.5% propyl gallate (Sigma-Aldrich, cat# 2370), 20mM Tris-HCl, pH 8.0 (invitrogen, cat# 15568-025).

Differential permeabilization was performed on HeLa cells transfected with the GFP-LBD constructs. Cells were fixed as described above 24 hours post transfection and permeabilized with 0.25% Triton X-100 or 0.015% digitonin for 5 min at RT following standard IF protocol with anti-GFP and anti-Lamin A/C antibodies.

### Antibodies

The following antibodies were used in this study: anti-H3K9me3 (Abcam, cat# ab8898), anti-Lamin A/C (Santa Cruz, cat# sc-376248), anti-Lamin B1 (Abcam, cat# ab16048), anti-GFP (Abcam, cat# ab290), anti-HA (Santa Cruz, cat# sc-7392).

### Image acquisition and analysis

Confocal immunofluorescent images were taken using a Leica TCS SP8 3X STED confocal microscope using 63x/1.40 oil objective. Images were acquired using HyD detectors in the standard mode with 100% gain. All images were taken with minimal laser power to avoid saturation. Confocal images were deconvoluted using Huygens Professional software (Scientific Volume Imaging, The Netherlands). Some images in the Online Supplemental material were captured with Nikon TE2000 or Leica SP8 confocal microscopes. Confocal channels shift alignment was performed using 0.1μm TetraSpeck fluorescent beads (Invitrogen, cat# T7279). Image analysis was performed using Image J software (National Institute of Health, USA). Line signal intensity profile plots were created using Plot Profile tool. Measurement of localization of the IF signal at the nuclear periphery was performed as a proportion of the signal at the nuclear periphery measured using a mask of the nuclear lamina, to total signal in the nucleus.

### Pulldown experiments

GFP-tagged PRR14 wild type/mutant Tantalus fragments were used as bait proteins to measure interaction with wild type/mutant HA-tagged PP2A B56alpha in transfected cells. ChromoTek GFP-Trap Magnetic Agarose affinity beads (GFP-Trap®_MA, cat# gtma-20) were used for immunoprecipitation of GFP-fusion proteins. The GFP-Trap®_MA is a GFP Nanobody/VHH coupled to magnetic agarose beads. HeLa cells (100 mm dishes) were co-transfected with pCMV6-AN-mGFP-PRR14 Tantalus constructs (or pCMV6-AN-mGFP) plus V245 pCEP-4HA B56alpha constructs using Lipofectamine 2000. Lysates were processed according to protocol provided by Chromotek. Briefly, cells were scraped in ice cold PBS and collected by centrifugation. After washing with cold PBS, cells were resuspended in 200 μl lysis buffer (10 mM Tris/Cl pH 7.5; 150 mM NaCl; 0.5 mM EDTA; 0.5% NP-40), plus protease inhibitors and 1 mM PMSF (Millipore Sigma cat# 10837091001). Tubes were kept on ice for 30 min, with pipetting, and centrifuged at 20,000 x g for 10 m at 4°C. The supernatant was diluted to 500 μl with dilution buffer (10 mM Tris/Cl pH 7.5; 150 mM NaCl; 0.5 mM EDTA). An aliquot of beads were washed in the dilution buffer, and finally resuspended in 500 μl in the same buffer. The diluted beads were added to the cleared lysate and were gently mixed for 1 hour at 4°C. The beads were collected, washed with dilution buffer, resuspend in 100 μl 2x SDS-sample buffer and heated for 10 min at 95°C to dissociate immunocomplexes. After removing the beads, samples were loaded on SDS-PAGE gels, and anti-GFP and anti-HA antibodies were used to detect GFP bait and HA-B56alpha prey proteins, respectively.

### Fluorescence recovery after photobleaching (FRAP)

FRAP imaging was performed on a Leica SP8 laser scanning confocal system equipped with a heating chamber, using HC PL APO 63x/1.4 NA CS2 oil immersion lens. HeLa cells were grown on 35mm MatTek dishes. Cells were transfected with either N-terminally tagged human PRR14 or pEGFP-D50 lamin A. FRAP was performed at 16-20 hours post-transfection, using 488 nm full laser power (65mW). Five bleach iterations were performed (2-3 s total, in 0.5 - 0.8-s interval). Recovery for GFP-PRR14 was measured for 58-60s at 2-s intervals. Recovery for Lamin A was measured for 5.75 - 6.0 min at 5-s intervals. Data were plotted graphically using GraphPad Prizm 8. EasyFrap software (Koulouras et al., 2018) was used to determine t1/2 and R-square values.

### Statistical analysis

Statistical analyses were performed with Graphpad PRISM 8.0.1 software (Graphpad Software, Inc.) using ANOVA one-way non-parametric (Kruskal-Wallis test) test with Dunn’s multiple comparison or unpaired non-parametric Student’s t-test (Mann-Whitney test).

## Acknowledgements

We thank Jonathan Chernoff, Jonathan Epstein, Johnathan Whetstine, and Cheryl Smith for discussions and comments on the manuscript. We thank Andrea Stout from the Penn CDB Microscopy Core, and the following Fox Chase Cancer Center Core Facilities and respective Managers for their assistance: Biological Imaging (Andrey Efimov), Cell Sorting (James Oesterling), Biostatistics and Bioinformatics (Michael Slifker), Cell Culture (Pam Nakajima), Molecular Modeling (Mark Andrake), and High Throughput Screening (Margret Einarson). We also thank Mikayla Fassler for assistance with site-directed mutagenesis and Margaret Baldini for assistance in imaging experiments. This work was supported by the following National Institutes of Health grants: R21 AG054829 (R.A.K), P30 CA006927 from the National Cancer Institute, and R35 HL140018 from the National Institutes of Health.

## Author contributions

R. A. Katz provided conceptualization, oversight, funding acquisition, and wrote the initial manuscript draft. A. Poleshko contributed to conceptualization, performed experiments, performed formal data analysis, provided resources, performed validation, prepared data, critically reviewed and edited the manuscript. K. L. Dunlevy, V. Medvedeva, J. E. Wilson, M. Hoque, T. Pellegrin, A. Maynard, M. E. Kremp, and J. S. Wasserman performed experiments. J. E. Wilson, and M. Hoque performed formal data analysis.

The authors declare no competing financial interests.

## Supplemental Information

**Figure S1.**
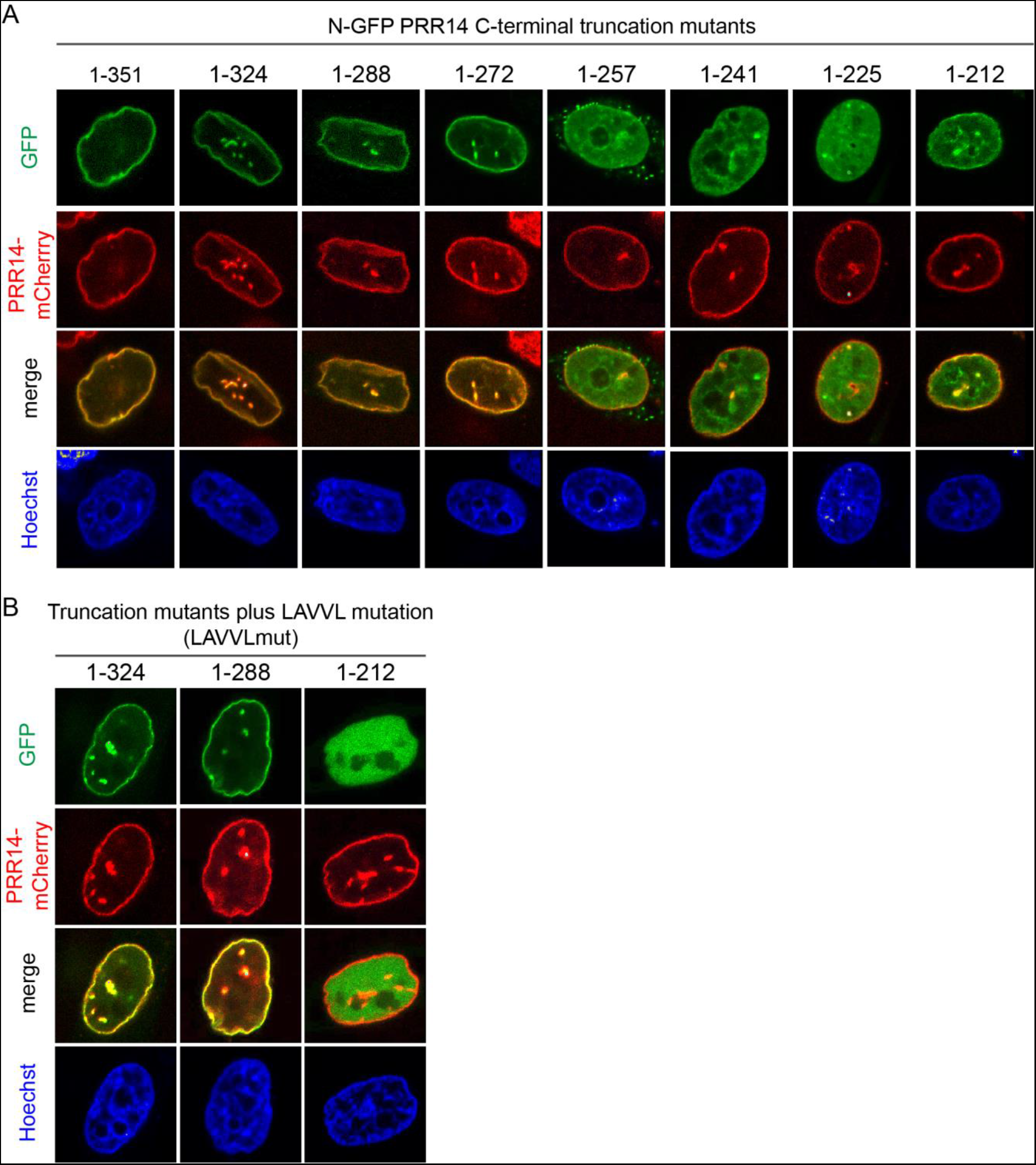
Deletion mapping of the PRR14 LBD. A series of PRR14 C-terminal truncations were created by introducing stop codons into the N-terminal GFP-tagged PRR14 reading frame. The end points of the C-terminal truncations are indicated. Representative confocal images of live HeLa cells stably expressing wild type mRFP-PRR14 transfected with **(A)** GFP-tagged truncation mutants. **(B)** composite mutants with substitutions in the LAVVL sequence required for heterochromatin binding (LAVVLmut). Counterstained with Hoechst.

**Figure S2.**
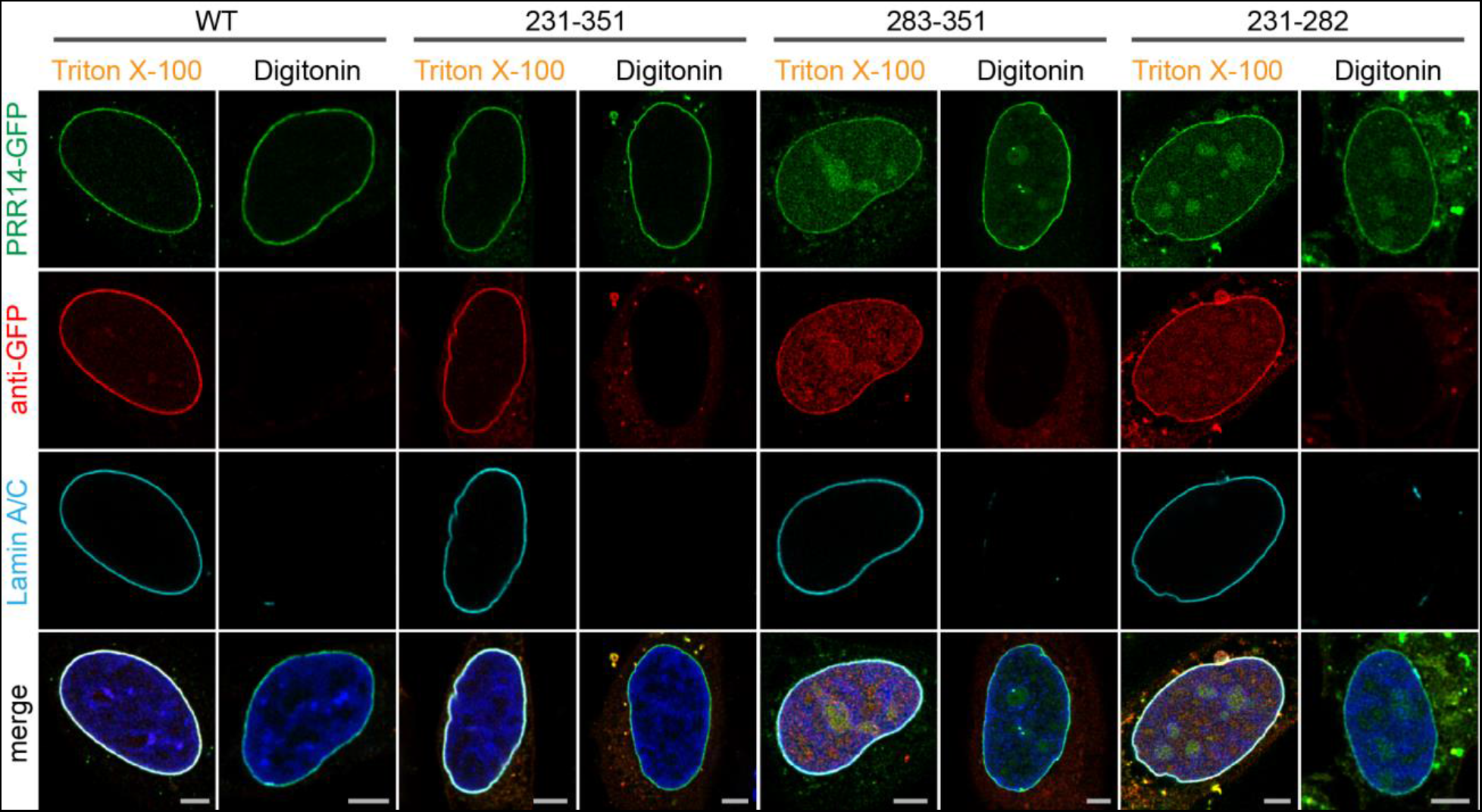
The PRR14 GFP-tagged 231-282 fragment localizes to the inner nuclear periphery. Representative confocal images of HeLa cells transfected with indicated GFP-PRR14 constructs and differentially permeabilized with Triton X-100 (plasma and nuclear membrane) or digitonin (plasma membrane only) to distinguish localization at the inner and outer nuclear periphery. Stained for GFP (red), Lamin A/C (cyan) and DAPI (blue). Scale bars: 5μm.

**Figure S3.**
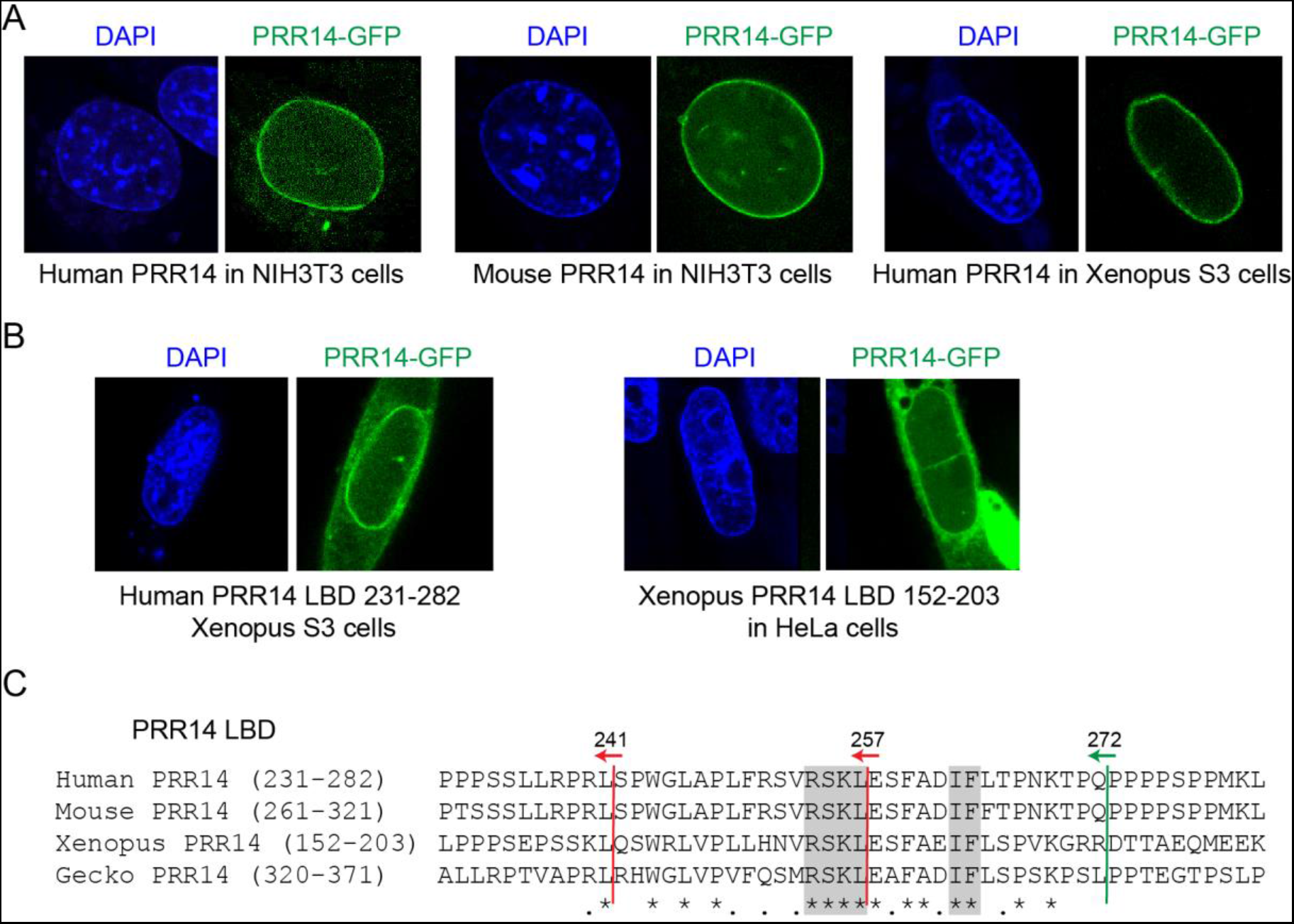
The mechanism for PRR14 nuclear lamina association is evolutionarily conserved. **(A)** GFP-tagged, full length human and mouse PRR14 proteins localize to the nuclear lamina in mouse cells. Human PRR14 localizes to the nuclear lamina in Xenopus cells. **(B)** GFP-tagged human PRR14 231-282 LBD localizes to the nuclear lamina in xenopus cells. GFP-tagged putative Xenopus PRR14 (see Table S1) 152-203 LBD localizes to the nuclear lamina in HeLa cells. **(C)** Manual alignment with no gaps of human and mouse, and putative xenopus and gecko LBDs. Amino acid identity (*) and similarity (.) are indicated. Conserved blocks chosen for analyses are indicated by shading. End points of relevant C-terminal deletions analyzed in Figure S1 are indicated. Green arrow indicates retention of nuclear lamina association and red arrows indicate loss of nuclear lamina localization. Deletion into the most conserved region of the human 231-282 LBD resulted in loss of nuclear lamina localization (Fig. S1).

**Figure S4.**
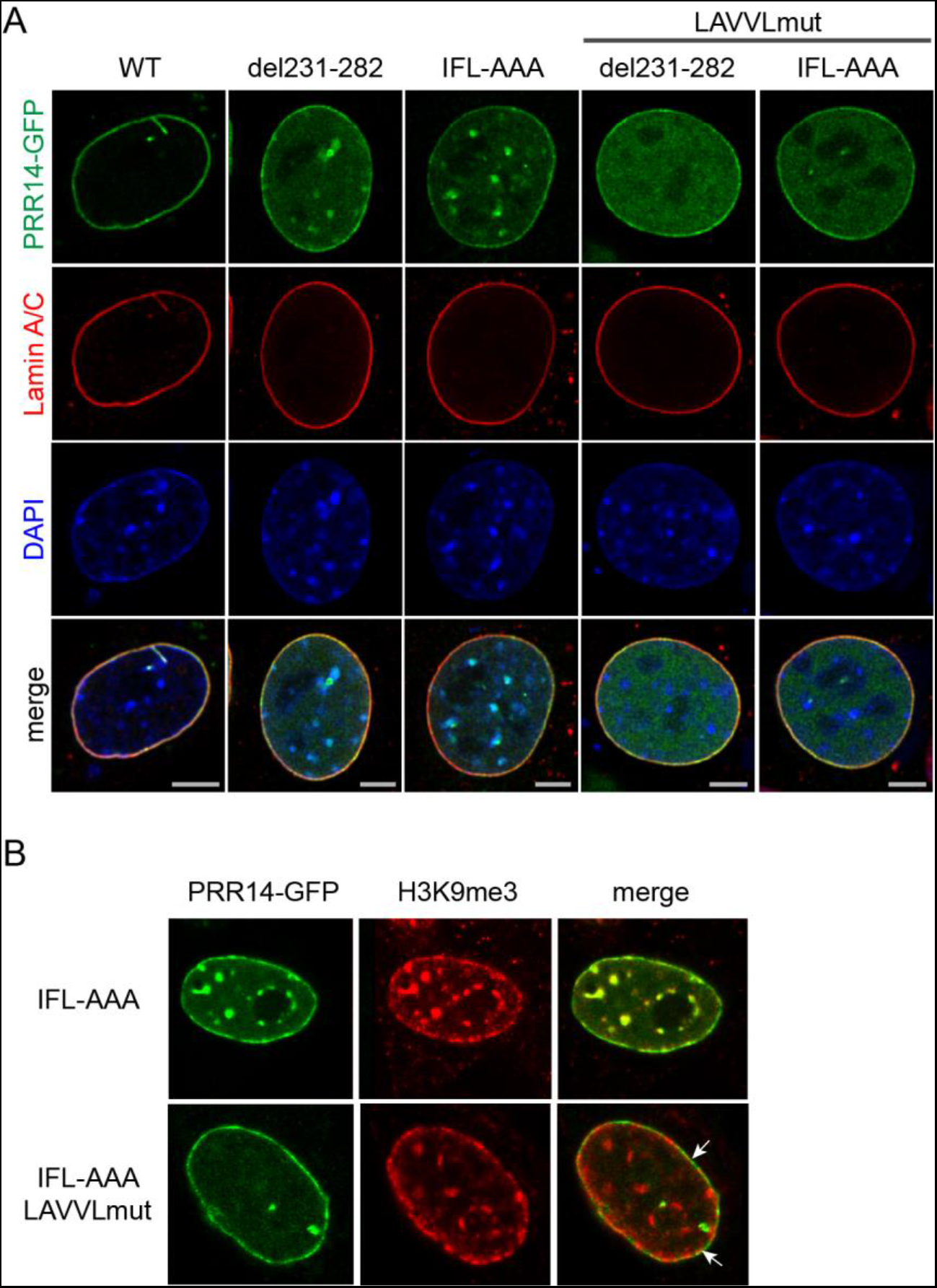
Composite mutations in the PRR14 231-282 LBD and HP1 binding site (LAVVLmut) result largely in nucleoplasmic localization. **(A)** Representative confocal images of murine C2C12 cells transfected with indicated PRR14-GFP constructs similar to experiment shown on figure 3B. The IFL-AAA and del231-282 constructs colocalization with chromocenters is observed. However, composite mutants incapable of HP1/heterochromatin binding (LAVVLmut) lack chromocenter localization and spread through nucleoplasm. Though, the composite mutants showed some localization to the nuclear periphery, suggesting residual nuclear lamina binding. **(B)** Representative confocal images of HeLa cells transfected with indicated IFL-AAA and composite IFL-AAA LAVVL-mutant, stained for H3K9me3. Colocalization of the IFL-AAA, but not the composite mutant protein with H3K9me3 heterochromatin is observed. Scale bars: 5μm.

**Figure S5.**
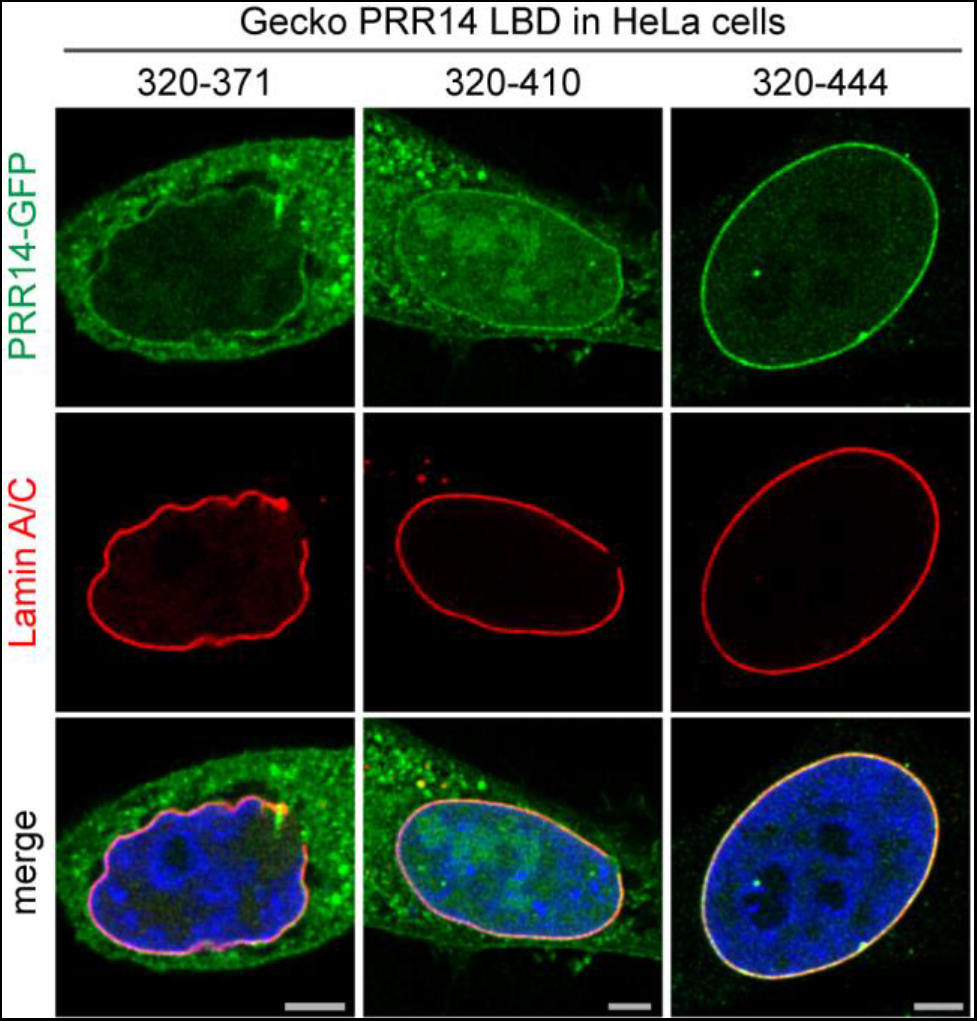
Mapping of the gecko PRR14 LBD. Representative confocal images of HeLa cells transfected with GFP-tagged gecko PRR14 320-371, 320-410, and 320-444 fragments (green), stained for Lamin A/C (red) and DAPI (blue). Scale bars: 5μm.

**Figure S6.**
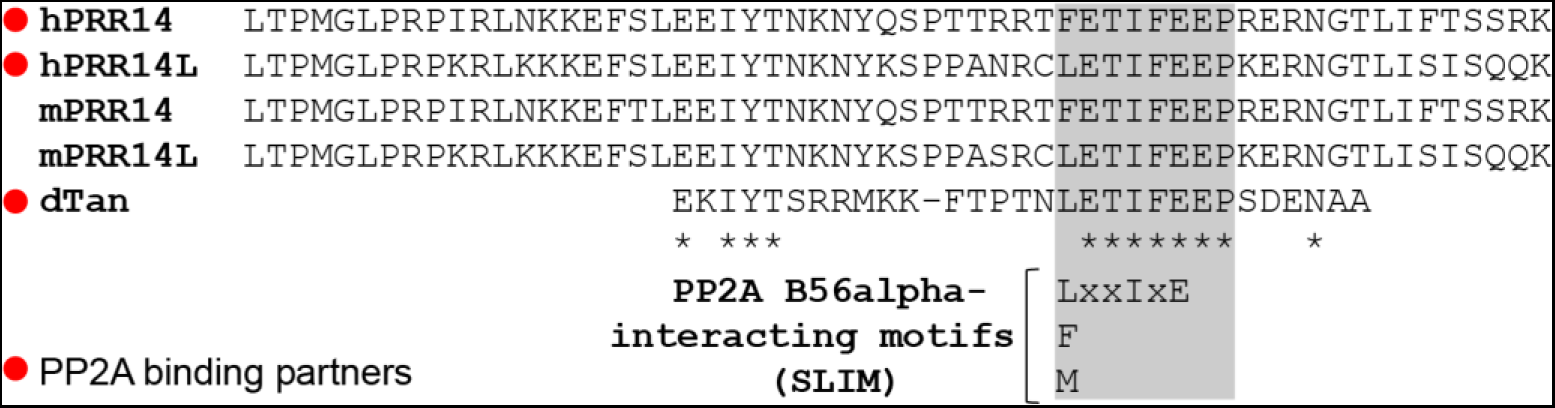
Alignment of Pfam Tantalus domains from mammalian PRR14, PRR14L and Drosophila Tantalus. Human and mouse PRR14 and PRR14L Tantalus domain sequences are shown, aligned to positions 145-174 of the 299 amino acid Drosophila Tantalus protein. The 145-174 Drosophila Tantalus region shown is only a subset of the 119-198 Drosophila Tantalus domain, as it was the only region showing significant homology. Human PRR14, PRR14L, and Drosophila Tantalus had been identified as PP2A interactors (see main text) implicating the conserved core sequences (shaded) in mediating PP2A binding. Subsequently, the L/F/MxxIxE motif was identified as a common recognition sequence for the B56alpha regulatory subunit of PP2A (see main text).

**Table S1.**
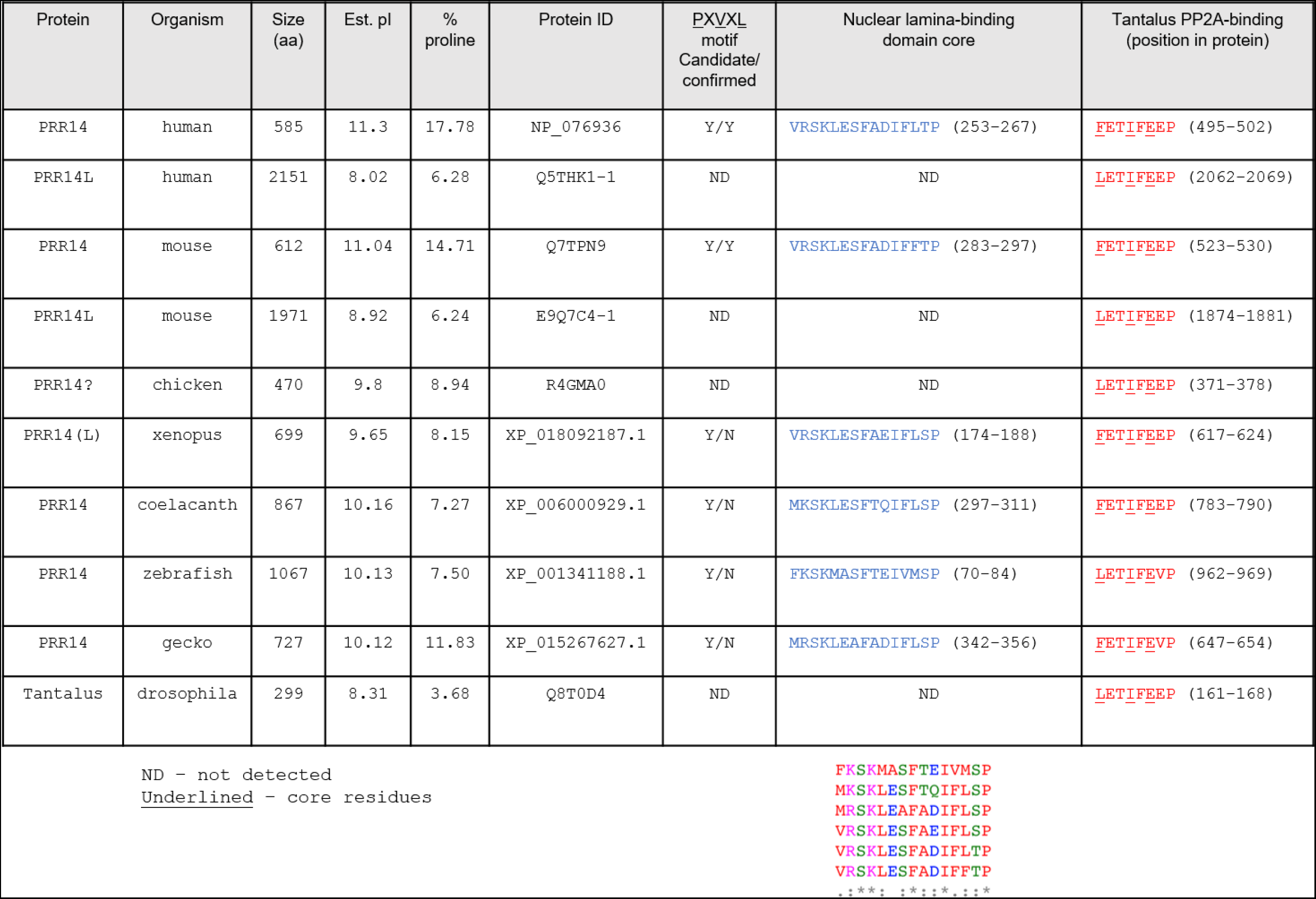
Comparison of features of mammalian PRR14 and the PRR14L paralog with candidate PRR14 genes in non-mammalian species. “ND” not detected, core residues underlined.

| Protein | Organism | Size (aa) | Est. pI | % proline | Protein ID | PXVXL motif Candidate/confirmed | Nuclear lamina-binding domain core | Tantalus PP2A-binding (position in protein) |
| --- | --- | --- | --- | --- | --- | --- | --- | --- |
| PRR14 | human | 585 | 11.3 | 17.78 | NP_076936 | Y/Y | VRSKLESFADIFLTP (253-267) | <u>FET</u><u>I</u><u>FE</u><u>EP</u> (495-502) |
| PRR14L | human | 2151 | 8.02 | 6.28 | Q5THK1-1 | ND | ND | <u>LET</u><u>I</u><u>FE</u><u>EP</u> (2062-2069) |
| PRR14 | mouse | 612 | 11.04 | 14.71 | Q7TPN9 | Y/Y | VRSKLESFADIFFTP (283-297) | <u>FET</u><u>I</u><u>FE</u><u>EP</u> (523-530) |
| PRR14L | mouse | 1971 | 8.92 | 6.24 | E9Q7C4-1 | ND | ND | <u>LET</u><u>I</u><u>FE</u><u>EP</u> (1874-1881) |
| PRR14? | chicken | 470 | 9.8 | 8.94 | R4GMA0 | ND | ND | <u>LET</u><u>I</u><u>FE</u><u>EP</u> (371-378) |
| PRR14 (L) | xenopus | 699 | 9.65 | 8.15 | XP_018092187.1 | Y/N | VRSKLESFAEIFLSP (174-188) | <u>FET</u><u>I</u><u>FE</u><u>EP</u> (617-624) |
| PRR14 | coelacanth | 867 | 10.16 | 7.27 | XP_006000929.1 | Y/N | MKSKLESFTQIFLSP (297-311) | <u>FET</u><u>I</u><u>FE</u><u>EP</u> (783-790) |
| PRR14 | zebrafish | 1067 | 10.13 | 7.50 | XP_001341188.1 | Y/N | FKSKMASFTEIVMSP (70-84) | <u>LET</u><u>I</u><u>FE</u><u>VP</u> (962-969) |
| PRR14 | gecko | 727 | 10.12 | 11.83 | XP_015267627.1 | Y/N | MRSKLEAFADIFLSP (342-356) | <u>FET</u><u>I</u><u>FE</u><u>VP</u> (647-654) |
| Tantalus | drosophila | 299 | 8.31 | 3.68 | Q8T0D4 | ND | ND | <u>LET</u><u>I</u><u>FE</u><u>EP</u> (161-168) |

ND - not detected Underlined - core residues
FKSKMASFTEIVMSP MKSKLESFTQIFLSP MRSKLEAFADIFLSP VRSKLESFAEIFLSP VRSKLESFADIFLTP VRSKLESFADIFFTP .:\*\*\*: :\*:~\*~\*~\*~\*~\*~\*

**Table S2.**
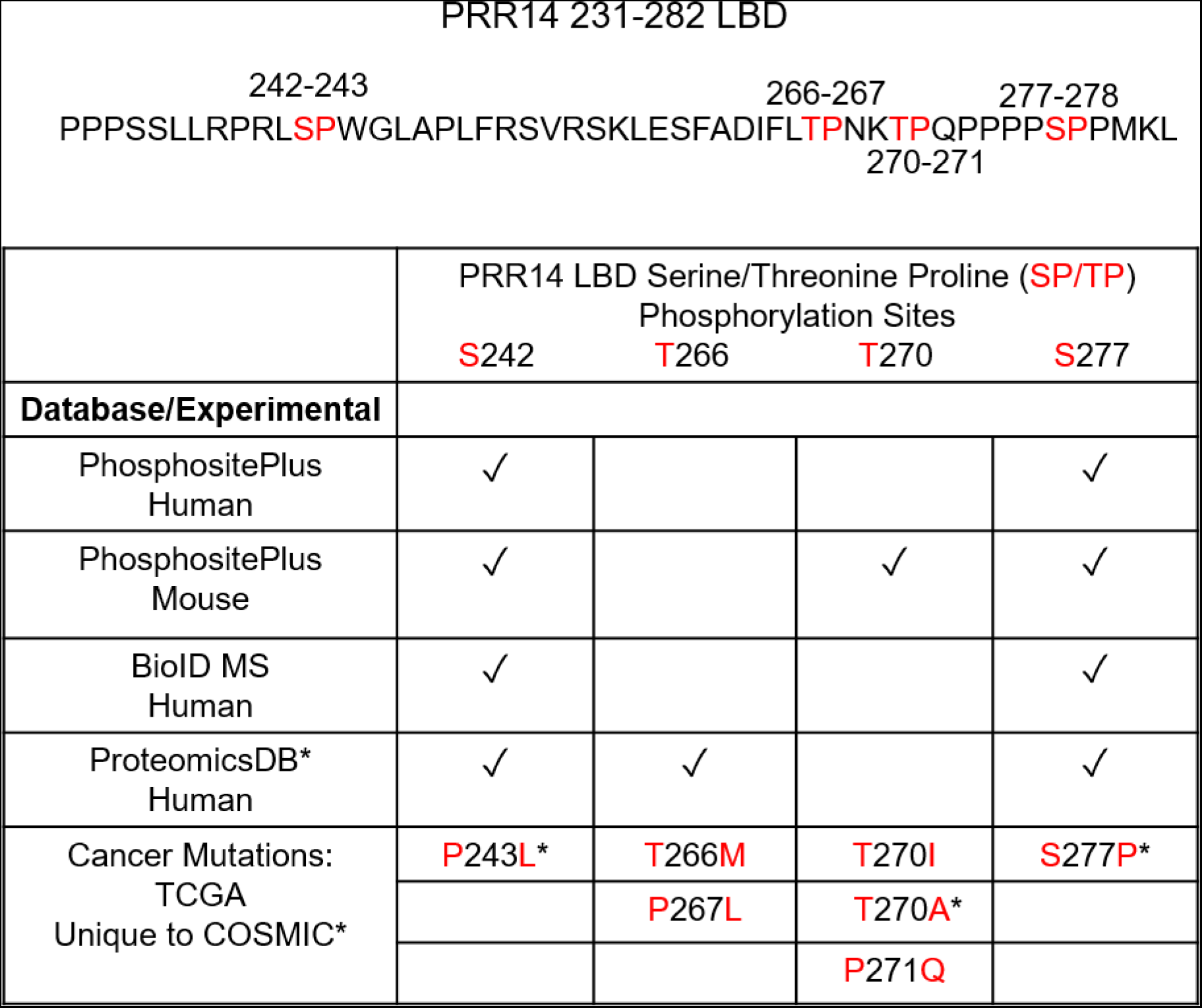
Summary of detection of PRR14 phosphopeptides in databases and our experimental work, BioID MS (mass spec). Top: Sequence of the PRR14 LBD showing the four candidate SP/TP phosphorylation sites.

Supplemental Information includes six supplemental figures, two supplemental tables, and two videos.

**Video S1. GFP-Lamin A fluorescence recovery at the nuclear periphery after photobleaching.** Represented movie of a HeLa cell transfected with Lamin A-GFP construct demonstrates slow Lamin A recovery at the nuclear periphery after photobleaching. Time stamp and scale bar are shown. Frames were captured in 5 s intervals.

**Video S2. GFP-PRR14 fluorescence recovery at the nuclear periphery after photobleaching.** Represented movie of a HeLa cell transfected with PRR14-GFP construct demonstrates rapid PRR14 recovery at the nuclear periphery after photobleaching. Time stamp and scale bar are shown. Frames were captured in 2 s intervals.

## References

Amendola, M., and B. van Steensel. 2014. Mechanisms and dynamics of nuclear lamina-genome interactions. Curr. Opin. Cell Biol. 28:61–68.

Aravind, L., and L.M. Iyer. 2012. The HARE-HTH and associated domains: novel modules in the coordination of epigenetic DNA and protein modifications. Cell Cycle. 11:119–131.

Bank, E.M., and Y. Gruenbaum. 2011. The nuclear lamina and heterochromatin: a complex relationship. Biochem. Soc. Trans. 39:1705–1709.

Becker, J.S., D. Nicetto, and K.S. Zaret. 2016. H3K9me3-dependent heterochromatin: Barrier to cell fate changes. Trends Genet. 32:29–41.

Buchwalter, A., J.M. Kaneshiro, and M.W. Hetzer. 2019. Coaching from the sidelines: the nuclear periphery in genome regulation. Nat. Rev. Genet. 20:39–50.

Bustin, M., and T. Misteli. 2016. Nongenetic functions of the genome. Science. 352:aad6933.

Butin-Israeli, V., S.A. Adam, A.E. Goldman, and R.D. Goldman. 2012. Nuclear lamin functions and disease. Trends Genet. 28:464–471.

Canzio, D., A. Larson, and G.J. Narlikar. 2014. Mechanisms of functional promiscuity by HP1 proteins. Trends Cell Biol. 24:377–386.

Chase, A., A. Pellagatti, S. Singh, J. Score, W.J. Tapper, F. Lin, Y. Hoade, C. Bryant, N. Trim, B.H. Yip, K. Zoi, C. Rasi, L.A. Forsberg, J.P. Dumanski, J. Boultwood, and N.C.P. Cross. 2019. PRR14L mutations are associated with chromosome 22 acquired uniparental disomy, age-related clonal hematopoiesis and myeloid neoplasia. Leukemia. 33:1184–1194.

Davidson, P.M., and J. Lammerding. 2014. Broken nuclei - lamins, nuclear mechanics, and disease. Trends Cell Biol. 24:247–256.

Dekker, J., A.S. Belmont, M. Guttman, V.O. Leshyk, J.T. Lis, S. Lomvardas, L.A. Mirny, C.C. O’Shea, P.J. Park, B. Ren, J.C.R. Politz, J. Shendure, S. Zhong, and D.N. Network. 2017. The 4D nucleome project. Nature. 549:219–226.

Dephoure, N., K.L. Gould, S.P. Gygi, and D.R. Kellogg. 2013. Mapping and analysis of phosphorylation sites: a quick guide for cell biologists. *Mol.Biol*. Cell. 24:535542.

Dephoure, N., C. Zhou, J. Villen, S.A. Beausoleil, C.E. Bakalarski, S.J. Elledge, and S.P. Gygi. 2008. A quantitative atlas of mitotic phosphorylation. Proc. Natl. Acad. Sci. U.S.A. 105:10762–10767.

Dietrich, B.H., J. Moore, M. Kyba, G. dosSantos, F. McCloskey, T.A. Milne, H.W. Brock, and H.M. Krause. 2001. Tantalus, a novel ASX-interacting protein with tissue-specific functions. Dev. Biol. 234:441–453.

Dittmer, T.A., and T. Misteli. 2011. The lamin protein family. Genome Biol. 12:222.

Eberhart, A., Y. Feodorova, C. Song, G. Wanner, E. Kiseleva, T. Furukawa, H. Kimura, G. Schotta, H. Leonhardt, B. Joffe, and I. Solovei. 2013. Epigenetics of eu- and heterochromatin in inverted and conventional nuclei from mouse retina. Chromosome Res. 21:535–554.

El-Gebali, S., J. Mistry, A. Bateman, S.R. Eddy, A. Luciani, S.C. Potter, M. Qureshi, L.J. Richardson, G.A. Salazar, A. Smart, E.L.L. Sonnhammer, L. Hirsh, L. Paladin, D. Piovesan, S.C.E. Tosatto, and R.D. Finn. 2019. The Pfam protein families database in 2019. Nucleic Acids Res. 47:D427–D432.

Giannios, I., E. Chatzantonaki, and S. Georgatos. 2017. Dynamics and Structure-Function Relationships of the Lamin B Receptor (LBR). PLoS One. 12:e0169626.

Glatter, T., A. Wepf, R. Aebersold, and M. Gstaiger. 2009. An integrated workflow for charting the human interaction proteome: insights into the PP2A system. Mol. Syst. Biol. 5:237.

Goldman, R.D., Y. Gruenbaum, R.D. Moir, D.K. Shumaker, and T.P. Spann. 2002. Nuclear lamins: building blocks of nuclear architecture. Genes Dev. 16:533–547.

Gonzalez-Sandoval, A., B.D. Towbin, V. Kalck, D.S. Cabianca, D. Gaidatzis, M.H. Hauer, L. Geng, L. Wang, T. Yang, X. Wang, K. Zhao, and S.M. Gasser. 2015. Perinuclear anchoring of H3K9-methylated chromatin stabilizes induced cell fate in *C. elegans* embryos. Cell. 163:1333–1347.

Gordon, M.R., B.D. Pope, J. Sima, and D.M. Gilbert. 2015. Many paths lead chromatin to the nuclear periphery. Bioessays. 37:862–866.

Gruenbaum, Y., and R. Foisner. 2015. Lamins: nuclear intermediate filament proteins with fundamental functions in nuclear mechanics and genome regulation. Annu. Rev. Biochem. 84:131–164.

Guelen, L., L. Pagie, E. Brasset, W. Meuleman, M.B. Faza, W. Talhout, B.H. Eussen, A. de Klein, L. Wessels, W. de Laat, and B. van Steensel. 2008. Domain organization of human chromosomes revealed by mapping of nuclear lamina interactions. Nature. 453:948–951.

Guruharsha, K.G., J.F. Rual, B. Zhai, J. Mintseris, P. Vaidya, N. Vaidya, C. Beekman, C. Wong, D.Y. Rhee, O. Cenaj, E. McKillip, S. Shah, M. Stapleton, K.H. Wan, C. Yu, B. Parsa, J.W. Carlson, X. Chen, B. Kapadia, K. VijayRaghavan, S.P. Gygi, S.E. Celniker, R.A. Obar, and S. Artavanis-Tsakonas. 2011. A protein complex network of Drosophila melanogaster. Cell. 147:690–703.

Harr, J.C., A. Gonzalez-Sandoval, and S.M. Gasser. 2016. Histones and histone modifications in perinuclear chromatin anchoring: from yeast to man. EMBO Rep. 17:139–155.

Hertz, E.P., T. Kruse, N.E. Davey, B. Lopez-Mendez, J.O. Sigurethsson, G. Montoya, J.V. Olsen, and J. Nilsson. 2016. A Conserved Motif Provides Binding Specificity to the PP2A-B56 Phosphatase. Mol. Cell. 63:686–695.

Herzog, F., A. Kahraman, D. Boehringer, R. Mak, A. Bracher, T. Walzthoeni, A. Leitner, M. Beck, F.U. Hartl, N. Ban, L. Malmstrom, and R. Aebersold. 2012. Structural probing of a protein phosphatase 2A network by chemical cross-linking and mass spectrometry. Science. 337:1348–1352.

Hetzer, M. 2010. The Nuclear Envelope. Cold Spring Harb. Perspect. Biol. 2.

Hochegger, H., S. Takeda, and T. Hunt. 2008. Cyclin-dependent kinases and cell-cycle transitions: does one fit all? Nat. Rev. Mol. Cell Biol. 9:910–916.

Hornbeck, P.V., B. Zhang, B. Murray, J.M. Kornhauser, V. Latham, and E. Skrzypek. 2015. PhosphoSitePlus, 2014: mutations, PTMs and recalibrations. Nucleic Acids Res. 43:D512–D520.

Kind, J., L. Pagie, H. Ortabozkoyun, S. Boyle, S.S. de Vries, H. Janssen, M. Amendola, L.D. Nolen, W.A. Bickmore, and B. van Steensel. 2013. Single-cell dynamics of genome-nuclear lamina interactions. Cell. 153:178–192.

Koulouras, G., A. Panagopoulos, M.A. Rapsomaniki, N.N. Giakoumakis, S. Taraviras, and Z. Lygerou. 2018. EasyFRAP-web: a web-based tool for the analysis of fluorescence recovery after photobleaching data. Nucleic Acids Res. 46:W467–W472.

Lechner, M.S., D.C. Schultz, D. Negorev, G.G. Maul, and F.J. Rauscher, 3rd. 2005. The mammalian heterochromatin protein 1 binds diverse nuclear proteins through a common motif that targets the chromoshadow domain. Biochem. Biophys. Res. Commun. 331:929–937.

Lemaitre, C., and W.A. Bickmore. 2015. Chromatin at the nuclear periphery and the regulation of genome functions. Histochem. Cell Biol. 144:111–122.

Lomberk, G., L. Wallrath, and R. Urrutia. 2006. The heterochromatin protein 1 family. Genome Biol. 7:228.

Machida, S., Y. Takizawa, M. Ishimaru, Y. Sugita, S. Sekine, J.I. Nakayama, M. Wolf, and H. Kurumizaka. 2018. Structural basis of heterochromatin formation by human HP1. Mol. Cell. 69:385–397 e388.

Meister, P., and A. Taddei. 2013. Building silent compartments at the nuclear periphery: a recurrent theme. Curr. Opin. Genet. Dev. 23:96–103.

Nozawa, R.S., K. Nagao, H.T. Masuda, O. Iwasaki, T. Hirota, N. Nozaki, H. Kimura, and C. Obuse. 2010. Human POGZ modulates dissociation of HP1alpha from mitotic chromosome arms through Aurora B activation. Nat. Cell Biol. 12:719–727.

Olins, A.L., G. Rhodes, D.B. Welch, M. Zwerger, and D.E. Olins. 2010. Lamin B receptor: multi-tasking at the nuclear envelope. Nucleus. 1:53–70.

Padeken, J., and P. Heun. 2014. Nucleolus and nuclear periphery: velcro for heterochromatin. Curr. Opin. Cell Biol. 28:54–60.

Peric-Hupkes, D., W. Meuleman, L. Pagie, S.W. Bruggeman, I. Solovei, W. Brugman, S. Graf, P. Flicek, R.M. Kerkhoven, M. van Lohuizen, M. Reinders, L. Wessels, and B. van Steensel. 2010. Molecular maps of the reorganization of genome-nuclear lamina interactions during differentiation. Mol. Cell. 38:603–613.

Poleshko, A., and R.A. Katz. 2014. Specifying peripheral heterochromatin during nuclear lamina reassembly. Nucleus. 5:32–39.

Poleshko, A., A.V. Kossenkov, N. Shalginskikh, A. Pecherskaya, M.B. Einarson, A. Marie Skalka, and R.A. Katz. 2014. Human factors and pathways essential for mediating epigenetic gene silencing. Epigenetics. 9:1280–1289.

Poleshko, A., K.M. Mansfield, C.C. Burlingame, M.D. Andrake, N.R. Shah, and R.A. Katz. 2013. The human protein PRR14 tethers heterochromatin to the nuclear lamina during interphase and mitotic exit. Cell Rep. 5:292–301.

Poleshko, A., P.P. Shah, M. Gupta, A. Babu, M.P. Morley, L.J. Manderfield, J.L. Ifkovits, D. Calderon, H. Aghajanian, J.E. Sierra-Pagan, Z. Sun, Q. Wang, L. Li, N.C. Dubois, E.E. Morrisey, M.A. Lazar, C.L. Smith, J.A. Epstein, and R. Jain. 2017. Genome-Nuclear Lamina Interactions Regulate Cardiac Stem Cell Lineage Restriction. Cell. 171:573–587 e514.

Politz, J.C., D. Scalzo, and M. Groudine. 2013. Something silent this way forms: the functional organization of the repressive nuclear compartment. Annu. Rev. Cell. Dev. Biol. 29:241–270.

Reddy, K.L., J.M. Zullo, E. Bertolino, and H. Singh. 2008. Transcriptional repression mediated by repositioning of genes to the nuclear lamina. Nature. 452:243–247.

Roux, K.J., D.I. Kim, M. Raida, and B. Burke. 2012. A promiscuous biotin ligase fusion protein identifies proximal and interacting proteins in mammalian cells. J. Cell Biol. 196:801–810.

Rual, J.F., K. Venkatesan, T. Hao, T. Hirozane-Kishikawa, A. Dricot, N. Li, G.F. Berriz, F.D. Gibbons, M. Dreze, N. Ayivi-Guedehoussou, N. Klitgord, C. Simon, M. Boxem, S. Milstein, J. Rosenberg, D.S. Goldberg, L.V. Zhang, S.L. Wong, G. Franklin, S. Li, J.S. Albala, J. Lim, C. Fraughton, E. Llamosas, S. Cevik, C. Bex, P. Lamesch, R.S. Sikorski, J. Vandenhaute, H.Y. Zoghbi, A. Smolyar, S. Bosak, R. Sequerra, L. Doucette-Stamm, M.E. Cusick, D.E. Hill, F.P. Roth, and M. Vidal. 2005. Towards a proteome-scale map of the human protein-protein interaction network. Nature. 437:1173–1178.

Scaffidi, P., and T. Misteli. 2005. Reversal of the cellular phenotype in the premature aging disease Hutchinson-Gilford progeria syndrome. Nat Med. 11:440–445.

Schmidt, T., P. Samaras, M. Frejno, S. Gessulat, M. Barnert, H. Kienegger, H. Krcmar, J. Schlegl, H.C. Ehrlich, S. Aiche, B. Kuster, and M. Wilhelm. 2017. ProteomicsDB. Nucleic Acids Res. 46:D1271–D1281.

Seeling, J.M., J.R. Miller, R. Gil, R.T. Moon, R. White, and D.M. Virshup. 1999. Regulation of beta-catenin signaling by the B56 subunit of protein phosphatase 2A. Science. 283:2089–2091.

Shevelyov, Y.Y., and D.I. Nurminsky. 2012. The nuclear lamina as a gene-silencing hub. Curr. Issues Mol. Biol. 14:27–38.

Shevelyov, Y.Y., and S.V. Ulianov. 2019. The Nuclear Lamina as an Organizer of Chromosome Architecture. Cells. 8.

Shi, Y. 2009. Serine/threonine phosphatases: mechanism through structure. Cell. 139:468–484.

Simon, D.N., and K.L. Wilson. 2013. Partners and post-translational modifications of nuclear lamins. Chromosoma. 122:13–31.

Solovei, I., A.S. Wang, K. Thanisch, C.S. Schmidt, S. Krebs, M. Zwerger, T.V. Cohen, D. Devys, R. Foisner, L. Peichl, H. Herrmann, H. Blum, D. Engelkamp, C.L. Stewart, H. Leonhardt, and B. Joffe. 2013. LBR and lamin A/C sequentially tether peripheral heterochromatin and inversely regulate differentiation. Cell. 152:584–598.

Stancheva, I., and E.C. Schirmer. 2014. Nuclear envelope: connecting structural genome organization to regulation of gene expression. Adv. Exp. Med. Biol. 773:209–244.

Stephens, A.D., E.J. Banigan, and J.F. Marko. 2019. Chromatin’s physical properties shape the nucleus and its functions. Curr Opin Cell Biol. 58:76–84.

Strom, A.R., A.V. Emelyanov, M. Mir, D.V. Fyodorov, X. Darzacq, and G.H. Karpen. 2017. Phase separation drives heterochromatin domain formation. Nature. 547:241–245.

Tatarakis, A., R. Behrouzi, and D. Moazed. 2017. Evolving models of heterochromatin: From foci to liquid droplets. Mol. Cell. 67:725–727.

Towbin, B.D., A. Gonzalez-Sandoval, and S.M. Gasser. 2013. Mechanisms of heterochromatin subnuclear localization. Trends Biochem. Sci. 38:356–363.

Ungricht, R., and U. Kutay. 2015. Establishment of NE asymmetry-targeting of membrane proteins to the inner nuclear membrane. Curr. Opin. Cell Biol. 34:135–141.

van Koningsbruggen, S., M. Gierlinski, P. Schofield, D. Martin, G.J. Barton, Y. Ariyurek, J.T. den Dunnen, and A.I. Lamond. 2010. High-resolution whole-genome sequencing reveals that specific chromatin domains from most human chromosomes associate with nucleoli. Mol. Biol. Cell. 21:3735–3748.

van Steensel, B., and A.S. Belmont. 2017. Lamina-associated domains: Links with chromosome architecture, heterochromatin, and gene repression. Cell. 169:780–791.

Vertii, A., J. Ou, J. Yu, A. Yan, H. Pages, H. Liu, L.J. Zhu, and P.D. Kaufman. 2019. Two contrasting classes of nucleolus-associated domains in mouse fibroblast heterochromatin. Genome Res. 29:1235–1249.

Virshup, D.M., and S. Shenolikar. 2009. From promiscuity to precision: protein phosphatases get a makeover. Mol. Cell. 33:537–545.

Wang, J., Z. Wang, T. Yu, H. Yang, D.M. Virshup, G.J. Kops, S.H. Lee, W. Zhou, X. Li, W. Xu, and Z. Rao. 2016. Crystal structure of a PP2A B56-BubR1 complex and its implications for PP2A substrate recruitment and localization. Protein Cell. 7:516–526.

Wong, X., and K.L. Reddy. 2015. Finding the middlemen in genome organization. Dev. Cell. 35:670–671.

Yanez-Cuna, J.O., and B. van Steensel. 2017. Genome-nuclear lamina interactions: from cell populations to single cells. Curr. Opin. Genet. Dev. 43:67–72.

Ye, Q., and H.J. Worman. 1996. Interaction between an integral protein of the nuclear envelope inner membrane and human chromodomain proteins homologous to Drosophila HP1. J. Biol. Chem. 271:14653–14656.

Zeng, W., A.R. Ball, Jr., and K. Yokomori. 2010. HP1: heterochromatin binding proteins working the genome. Epigenetics. 5:287–292.

Zhou, V.W., A. Goren, and B.E. Bernstein. 2011. Charting histone modifications and the functional organization of mammalian genomes. Nat. Rev. Genet. 12:7–18.

Zullo, J.M., I.A. Demarco, R. Pique-Regi, D.J. Gaffney, C.B. Epstein, C.J. Spooner, T.R. Luperchio, B.E. Bernstein, J.K. Pritchard, K.L. Reddy, and H. Singh. 2012. DNA sequence-dependent compartmentalization and silencing of chromatin at the nuclear lamina. Cell. 149:1474–1487.

